# Targeting the translation initiation complex in therapy-resistant and metastatic melanoma

**DOI:** 10.1101/2024.11.20.624566

**Authors:** Yongmei Feng, Ximena Diaz-Olea, Xiurong Ke, Mariia Radaeva, Mehdi Amiri, Hyunjeong Joo, Maryam Jafari, Hyungsoo Kim, Aniruddha J. Deshpande, Steven Olson, Predrag Jovanovic, Yangqi Su, Aagam Shah, Ian Pass, Qiyun Deng, Ikrame Lazar, Rabi Murad, Alfredo Molinolo, Mark Faries, Saugato Rahman Dhruba, Tiangen Chang, Omid Hamid, Eduard Sergienko, Wei Li, Jessie Villanueva, Simon Knott, Ivan Topisirovic, Michael Jackson, Eytan Ruppin, Cristina Ferrone, Nahum Sonenberg, Artem Cherkasov, Ze’ev A. Ronai

## Abstract

Expression of components of the translation initiation complex (eIF4F) is frequently elevated in cancer, resulting in enhanced synthesis of oncogenic proteins. We thus set out to limit eIF4F pro-oncogenic activity, a notable challenge given its essential role in normal tissues. CRISPR-Cas9-based functional screen using tiling sgRNAs identified the eIF4G1 MA3 domain, a subunit of the eIF4F complex, as a target for developing small molecule inhibitors. Combination of structure-guided *in silico* modeling and chemical library screening led to the identification of small molecule candidates M19 and its analog M19-6 that binds to the MA3 domain of eIF4G1 and disrupts eIF4F complex. M19-6 treatment reprograms the melanoma translatome, limiting synthesis of factors that promote cell proliferation and neoplastic growth, as well as reducing translation of mRNAs encoding ferroptosis suppressors. Whole genome CRISPR screen indentified ferroptosis activators to augment M19-6 activity, which was confirmed in cultured melanoma cells. M19-6 alleviates melanoma resistance to BRAF and MEK inhibitors, while eliciting anti-tumor and anti-metastatic effects in preclinical mouse models. Among several biomarkers found in M19-6 sensitive cell lines, THBS1 and TGFβI expression were elevated in patients who are non-responders to PD-1 therapy as in patients with metastasis. Our studies identify M19-6 as a therapeutic candidate, offering a novel insights into targeting the eIF4F complex to overcome melanoma resistance to therapy and metastasis.

**Significance:** We identify M19-6 as a small molecule that disrupts the eIF4F, translation initiation complex, by targeting the MA3 domain of eIF4G1, resulting in elimination of melanoma cells in culture, overcoming therapy resistance while inhibiting melanoma growth and metastasis *in vivo*. As M19-6 causes minimal toxicity to melanocytes, it offers a therapeutic modality for melanoma.

## Background

Malignant melanoma is a rapidly metastasizing tumor that has served as a flagship model for cancer immunotherapy (1), although challenged by emerging resistance, which limits therapy effectiveness (2). Among factors proposed to hinder immunotherapy and induce drug resistance are components of the eIF4F complex (3-5). However, a significant challenge in targeting the activity of eIF4F complex in tumors is that this complex is essential for the viability of all cells.

Assembly of the eIF4F complex is the rate-limiting step in translation initiation (6). Although eIF4F activity is required for cap-dependent translation of all cytoplasmic mRNAs, there is a notable difference in requirements for eIF4F by housekeeping mRNAs (which are efficiently translated even when eIF4F complex activity or levels are low) versus cancer-promoting mRNAs (which require high eIF4F levels for their optimal translation) (7,8). Accordingly, eIF4F activity is frequently higher in malignant than in normal cells (7, 9,10), providing a clinically exploitable therapeutic window (7, 9). Elevated eIF4F levels, commonly seen in tumors, including melanoma (11, 12), correlate with poor prognosis (4, 11), and increased eIF4F activity is linked to mTORC1 hyperactivation and 4E-BP inactivation (13). As the major eIF4G paralogue, eIF4G1 is frequently overexpressed in malignancies, where it reportedly contributes to tumorigenesis and cancer progression, in part by regulating the DNA damage response and cell cycle activity (13-16).

Among small molecules that disrupt the eIF4F complex, we previously identified SBI-756, which binds to eIF4G1 and disrupts the eIF4F by targeting eIF4G1 independently of cellular 4E-BP status (5). This mechanism allows SBI-756 to act regardless of the eIF4E/4E-BP ratio in the cell, which is often unfavorable thus impeding current eIF4F-targeting modalities including mTOR inhibitors (13). Given that several oncogenes and tumor suppressors, including Myc and p53 (4,17), converge on the eIF4F complex, targeting this complex holds promise in overcoming challenges associated with tumor heterogeneity, metastasis, and drug resistance (4,5,14,18). eIF4F inhibitors currently being tested in preclinical studies and clinical trials include 4EGI-1, which binds to eIF4E and prevents its association with eIF4G in a 4E-BP-dependent manner; the eIF4A inhibitors hippuristanol and rocaglates (e.g., silvestrol); and cap analogs and mimetics that inhibit eIF4E binding to the 5’ mRNA cap (4, 19). Among the more recently developed eIF4F complex inhibitors is eFT226 (zotatifin), which inhibits eIF4A activity (20) and has demonstrated effectiveness against *Kras*-mutant lung cancer cells and triple-negative breast cancer (21, 22). eFT226 has now reached clinical evaluation. Finally, a second-generation, more potent eIF4A inhibitor, MG-002, has recently been developed (23).

We recently demonstrated that SBI-756 inhibits melanomagenesis in several mouse models of human and mouse cell xenografts, including the genetic *Nras*^*Q61K*^*::Ink4a*^*–/–*^ melanoma model (24). Independent studies have shown that SBI-756 is effective in a B-cell lymphoma model when combined with the BCL-2 inhibitor venetoclax (25). Recent studies have also revealed that SBI-756 may promote anti-tumor immunity and attenuate the growth of pancreatic tumors in preclinical models (26). SBI-756 treatment also promoted a gene expression signature resembling that caused by genetic ablation of eIF4G1 (14). Analysis of genetic changes in melanoma cells resistant to SBI-756 identified several genes associated with the cell cycle and DNA repair, hallmarks of eIF4G1 inhibition (14). Nonetheless, suboptimal biophysical properties of SBI-756 have limited its further development, prompting the creation of a new generation of eIF4G1 inhibitors.

Here, we identify the MA3 domain of eIF4G1 as the target of SBI-756 and categorize a new class of small molecules that directly target the MA3 domain. Among the selected small molecules, M19 and its analog M19-6 disrupt the eIF4F complex and interfere with eIF4A/eIF4G1 interaction. Ribo-Seq analysis identified that M19-6 limited the translation of mRNAs implicated in cell cycle, protein synthesis and ferroptosis. A whole genome CRISPR screen to identify pathways that can complement M19-6 activity unraveled that ferroptosis promoting factors increase M19-6 toxicity towards melanoma cells. Among biomarkers that identify M19-6-sensitive melanoma cell lines are THBS1 and TGFβI, which were also elevated in non-responders to PD-1 therapy. *In vivo*, M19-6 effectively attenuated melanoma growth and metastasis. The nature of M19-6 provides a new therapeutic modality to reduce the growth and metastatic spread of melanoma.

## Results

### Identification of the eIF4G1 MA3 domain as an SBI-756 target

To identify the eIF4G1 domain that binds SBI-756, we transduced the A375 and UACC903 human melanoma lines, both of which stably expressed Cas9 (see Methods for details), with a tiling sgRNA library consisting of 947 guides to target the eIF4G1 gene. After selecting sgRNA-expressing cells, we treated them with increasing SBI-756 concentrations and then harvested viable cells for DNA extraction and sequencing to identify sgRNAs conferring SBI-756 resistance. That analysis identified two gRNAs targeting sequences either within (gRNA-235) or proximal to (gRNA-622) the eIF4G1 MA3 domain (27) (Figure 1a, Supplementary Figure 1a), suggesting that this domain is required for SBI-756 targeting. To verify that these sgRNAs conferred SBI-756 resistance, we transduced A375 melanoma cells with either gRNA-235 or gRNA-622 and selected individual clones that became relatively more resistant to SBI-756 than controls. One clone transduced with gRNA-235 exhibited a 14-base pair deletion within the MA3 domain, resulting in early termination of eIF4G1 mRNA and protein. In contrast, the clone transduced with gRNA-622 exhibited a 12-base pair deletion within a domain proximal to MA3, which did not promote early termination of eIF4G1 transcripts (Supplementary Figure 1b). Previous studies suggest that the MA3 domain is important for eIF4G1/eIF4A interaction (28,29). These findings are consistent with our analysis performed in *S. Cerevisiae* transduced with an eIF4F gene set library and aimed at identifying components of the translation initiation machinery that confer SBI-756 resistance. That search revealed that expression of truncated forms of eIF4G1 and eIF4A1 genes (TIF1) is sufficient to confer SBI-756 resistance (Supplementary Figure 1c).

**Figure 1.**
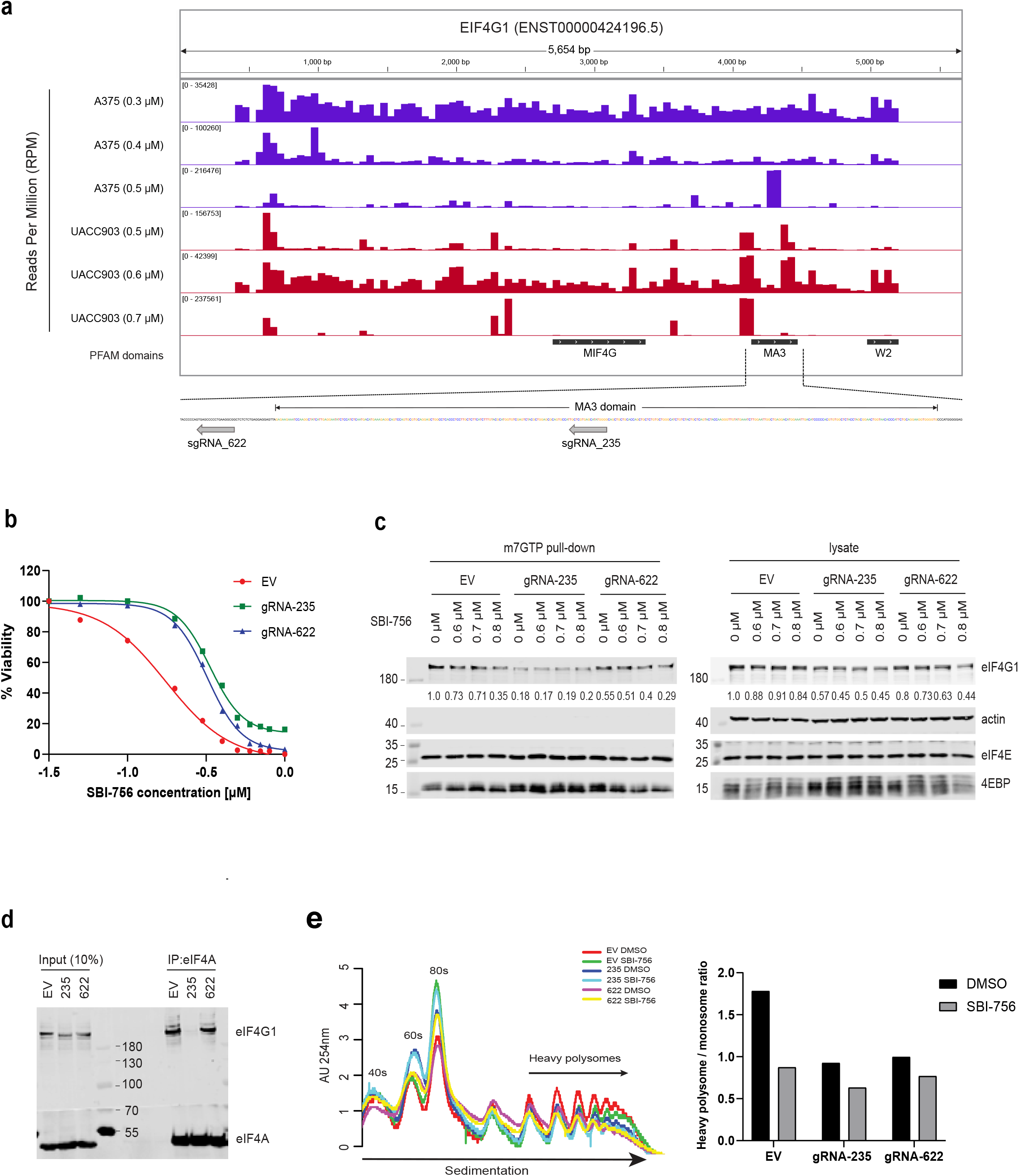
Identifying MA3 domain as SBI-756 target on eIF4G1. **CRISPR screen identifies eIF4G1 domains targeted by SBI-756**. A CRISPR library of sgRNAs prepared against different eIF4G1 gene domains was used to transduce A375 and UACC903 human melanoma cells, which were then treated with puromycin to select for sgRNA-expressing cells and with increasing SBI-756 concentrations. **(a)** Genomic DNA from cells was PCR-amplified and Sanger-sequenced. Read counts were mapped to each sgRNA. Two enriched sgRNAs (gRNA-235 and gRNA-622) were mapped to the eIF4G1 MA3 domain. gRNA-235 falls within the MA3 domain and results in early termination (aa 1322 of 1606) and thus excludes the W2 domain, which reportedly interacts with eIF4A, from this transcript. gRNA-622 is located proximal (upstream) of the MA3 domain. **(b)** MA3 mutant melanoma cells are more resistant to SBI-756 than parental MA3 domain WT melanoma cells. gRNA-235 and gRNA-622 melanoma cells obtained as described in Methods were treated with indicated concentrations of SBI-756, followed by the analysis of their viability (72 h later) using CellTier-Glo. **(c)** M^7^GTP pull-down assay. Proteins prepared from gRNA-235, gRNA-622 A375 melanoma cells that were treated with indicated SBI-756 concentrations were incubated with m^7^GTP-agarose beads to capture the eIF4F complex. Indicated proteins were detected by Western blot analysis. Each protein band was quantified by ImageJ, normalized to eIF4E levels as indicated. Right panel shows the total cell lysate. Each protein band was quantified by ImageJ, normalized to actin levels as indicated. **(d)** Proteins prepared from EV, gRNA-235 and gRNA-622 A375 cells were immunoprecipitated with eIF4A antibody and detected with the indicated antibodies. **(e)** Polysome profiling of human melanoma A375 cells harboring WT eIF4G1 or eIF4G1 with a mutant MA3 domain (gRNA-235 and gRNA-622), prior to and after SBI-756 treatment. Three replicates are shown, as is the polysome/monosome ratio (right panels).

We next assessed viability of single clones expressing either gRNA-235 or gRNA-622 after SBI-756 treatment. Compared to control (EV) sgRNA-transduced cells, melanoma cells expressing either gRNA-235 or gRNA-622 exhibited increased SBI-756 resistance (Figure 1b). gRNA-235-transduced melanoma cells also showed decreased proliferation relative to EV- or gRNA-622-transduced cells (Supplementary Figure 1d).

Since SBI-756 disrupts eIF4F complex assembly (6), we asked whether melanoma cells expressing either gRNA-235 or gRNA-622 alter association of eIF4F subunits. Using pull-down analysis with the methyl-^7^GTP cap analogue to capture complex components, we observed that eIF4F complex levels were lower in melanoma cells expressing gRNA-235, as indicated by a notable decrease in eIF4G1 and increased abundance of 4E-BP1 in pulled-downs, as compared to EV controls or gRNA-622-expressing cells (Figure 1c). Conversely, SBI-756 treatment disrupted eIF4F complex formation in EV cells, but not in gRNA-235-expressing cells. Interestingly, gRNA-622-expressing cells did not exhibit eIF4F complex dissociation (Figure 1c), suggesting that a different mechanism underlies melanoma resistance to SBI-756 in this context. Given that the MA3 domain is implicated in eIF4A/eIF4G1 interaction, we investigated whether this interaction was disrupted in melanoma cells expressing mutant eIF4G1 forms. eIF4A/eIF4G1 interaction was disrupted by SBI-756 in melanoma cells expressing gRNA-235 but not gRNA-622 (Figure 1d). This further suggests that distinct mechanisms mediate effects of gRNA-235 and gRNA-622 on melanoma cell response to SBI-756.

To assess potential impact of MA3 domain disruption on overall translation activity, we performed polysome profiling (30,31) of individual gRNA-235 or gRNA-622-expressing cells, relative to control, non-MA3 domain targeted cells (Figure 1e, left panel). Indeed, polysome profiles in vehicle treated gRNA-235- or gRNA-622-expressing cells resembled those observed in non-MA3 domain targeted cells treated with SBI-756 (Figure 1e, left panel). Moreover, relative to vehicle controls, SBI-756 treatment depleted polysomes while increasing monosome peak in control, MA3 domain intact cells, but not those expressing gRNA-235 or gRNA-622 (Figure 1e, left panel). These observations, confirmed by quantifying heavy polysome/monosome ratios (Figure 1e, right panel), suggest that MA3 mutations generated in our CRISPR screen are likely to limit the effect of SBI-756 on the eIF4F complex and thus translation initiation. Collectively, both gRNA-622- and gRNA-235-expressing cells have equally limited polysomes, substantiating the importance of protein synthesis in resistance to SBI-756.

### An in silico screen identifies small molecules binding to the MA3 domain

To identify small-molecule inhibitors targeting MA3 domain, we conducted an *in silico* screen based on the human eIF4GI crystal structure (PDB:1UG3) of a surface binding site in the protein’s C-terminus (32). That site included amino acid residues Ala1292, Ile1293, Glu1343, Thr1346, Gln1350, Leu1384, and Lys1386 within the MA3 domain, which is predicted to be essential for drug binding. This selection was informed by blind docking experiments with SBI-756 and assessment of druggability of the MA3 domain surface. Molecular dynamics simulation (30ns, using the Desmond suite from Schrodinger; (33)) also revealed that SBI-756 formed stable interactions with these residues (Figure 2a). Druggability of the selected pocket was strongly suggested by both the depth and hydrophobic characteristics of the binding cavity and stability of these interactions (Figure 2a).

**Figure 2.**
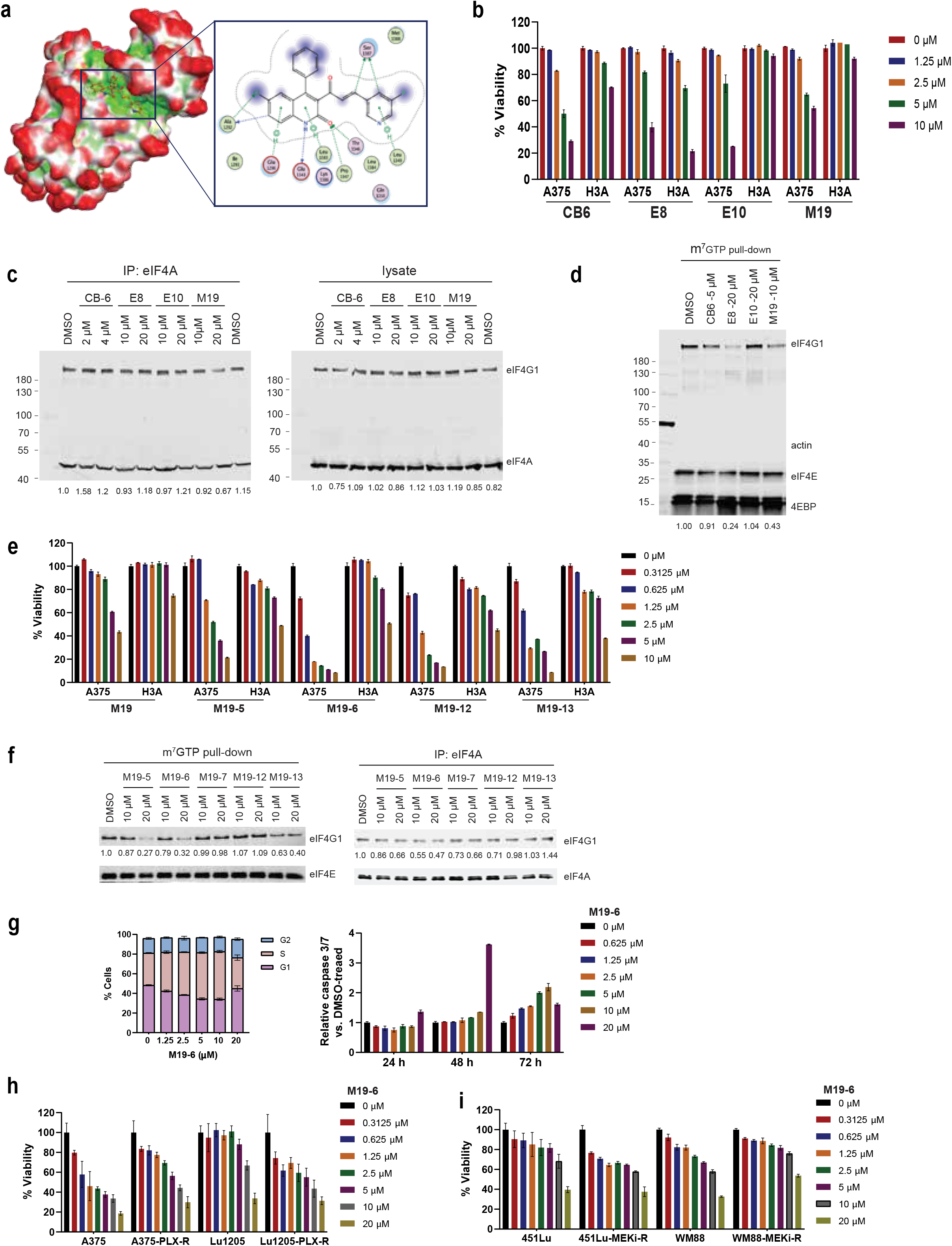
In Silico screen to identify MA3 domain bound small molecules. **(a)** The proposed binding mode of SBI-756 to the surface of the eIF4G1 MA3 domain. Left, high-view of SBI-756 binding to the MA3 domain (PDB: 1UG3). Green areas, hydrophobic patches; red areas, solvent exposed. Right, ligand-protein interactions of SBI-756 inside the proposed pocket. Blue shading, solvent-exposed; lines with arene symbols, hydrophobic interactions; arrowed lines, hydrogen bonds. Bonds shown were maintained throughout the 30ns molecular dynamics simulation. **(b)** A375 and H3A cells were seeded in 96-well plates and treated with different concentrations of indicated compounds. Cell viability was measured after 72 h. **(c)** A375 cells were treated as indicated. After 24 h, cells were harvested followed by protein extraction, immunoprecipitation and Western blot analysis with the indicated antibodies. Each protein band was quantified by ImageJ, normalized to eIF4A levels as indicated. **(d)** Proteins prepared from A375 cells treated as indicated were incubated with m^7^GTP-agarose beads to capture the eIF4F complex. Indicated proteins were detected by Western blot analysis. Each protein band was quantified by ImageJ, normalized to eIF4E levels as indicated. **(e)** A375 and H3A cells were seeded in 96-well or 384-well Corning spheroid microplates. After 24 h, indicated compounds were added to respective wells and cell viability was measured after 4 days using CellTiter-Glo 3D. **(f)** M^7^GTP pull-down assay and immunoprecipitation assay. Left, proteins prepared from A375 cells treated with the indicated compounds were incubated with m^7^GTP-agarose beads to capture the eIF4F complex. Indicated proteins were detected by Western blot analysis. Each protein band was quantified by ImageJ, normalized to eIF4E levels as indicated below the blots. Right, proteins prepared from A375 cells treated with the indicated compounds were incubated with anti-eIF4A antibody followed by Western blot analysis. Each protein band was quantified by ImageJ, normalized to eIF4A levels as indicated. (**g**) Left, A375 cells were treated with indicated concentrations of M19-6. After 24h, cells were collected and fixed followed by FACS analysis. Right, A375 cells were treated with indicated concentrations of M19-6. Apoptosis was measured using caspase-Glo 3/7 kit. **(h)** and **(i)** BRAF inhibitor-resistant and MEK inhibitor-resistant cell lines were grown as spheroids and treated with M19-6. Cell viability was measured after 4 days.

Next, to identify small-molecule eIF4G1 inhibitors, we performed virtual screening of the MA3 binding pocket by screening a library of 10 million drug-like compounds through molecular docking followed by filtering steps (34). The initial docking phase identified 54,500 compounds exhibiting more favorable docking scores than the benchmark compound SBI-756. From this pool, we selected a chemically diverse subset of 500 compounds using filtration tools described in the Methods section. Final selection was carried out to exclude compounds with undesirable properties such as toxicity, promiscuity, and low solubility, resulting in 150 compounds, which were then clustered based on chemical diversity to yield 67 potential MA3 domain inhibitors (Supplementary Figure 2a).

### Assessment of lead MA3-interacting compounds in melanoma

Among the 67 potential compounds targeting the MA3 domain of eIF4G1, described above, 63 were available and purchased from ChemBridge (CB), Enamine (E), and Molport (M). We assessed all 63 for (i) toxicity to melanoma and other tumor lines, (ii) effect on melanoma lines relative to non-transformed melanocytes, and (iii) impact on gRNA-235-transduced melanoma cell lines. Initially, we tested compounds for toxicity against six melanoma lines representing diverse genetic mutations, as well as two prostate cancer cell lines. Of the 63 compounds evaluated, 4 (CB6, E8, E10, and M19) were selected for further analyses (Supplementary Figure 2b), and among them, E10 and M19 exhibited selectivity for melanoma cells relative to the immortalized melanocyte line H3A (Figure 2b). Relative to EV-transduced controls, compound M19 was less toxic to A375 cells expressing gRNA-235 (Supplementary Figure 2c). Decreasing serum levels further sensitized A375 melanoma cells to M19 treatment (Supplementary Figure 2d). Finally, among selected compounds, M19 caused the most potent disruption of both eIF4G1:eIF4A binding and eIF4F complex assembly, while eliciting the lowest toxicity (Figures 2c, 2d). Collectively, these data motivated us to select M19 for further experimental validation.

Next, we performed a similarity search using the M19 scaffold as the basis to identify analogs with potentially greater efficacy. Using the ZINC20 library, we retrieved 3000 compounds with a Tanimoto similarity score above 0.65. Docking scores of analogs were compared against the parental M19 compound identified and subsequently acquired 13 M19 analogs, which were predicted to have enhanced binding affinity to the MA3 domain (see methods for details). Of those, four (M19-5, M19-6, M19-12, and M19-13) demonstrated enhanced potency relative to M19 itself (Figure 2e, Supplementary Figure 2e). Of these, M19-6 was the most potent compound both in colony-forming (CFE, Supplementary Figure 2f), and in promoting eIF4F complex dissociation (Figure 2f).

Analysis of mRNA translation in the presence of M19-6 or other protein synthesis inhibitors revealed dose-dependent inhibition by M19-6. In this assay, M19-6 was more potent than the commercial inhibitor 4EGI-1, which blocks eIF4E-eIF4G1 binding and is known to be toxic (Supplementary Figure 2g).

We next assessed the effects of M19-6 in cancer cell lines grown in 2D or 3D (spheroid) cultures. Both analyses revealed selective effectiveness of M19-6 against A375, WM793, SKMEL5, H1693, the breast cancer ZR75-1 cells, as well as against BRAF inhibitor-resistant A375 cells, but not against the IGR37, IGR39, SKMEL28, WM3629 or MeWO cells (Supplementary Figure 2h). To determine whether NRAS or BRAF mutations might confer greater sensitivity to M19-6, we treated melanoma cells harboring these mutations with M19-6. NRAS-mutant melanoma cells exhibited greater sensitivity to M19-6 compared with BRAFmutant melanoma cells (Supplementary Figure 2i). Comparable to changes seen in melanoma cells treated with SBI-756 and the parental M19 compound, M19-6 induced more robust S1 arrest and apoptotic effects in melanoma cells, effects that exhibited time dependency (Figure 2g). Spheroids formed from different BRAF or MEK inhibitor-resistant melanoma lines were also M19-6 sensitive (Figure 2h, 2i).

### M19 treatment of melanoma cells perturbs gene expression

To identify possible changes in gene expression following M19 treatment, we performed RNA-Seq using A375 melanoma cells treated with either vehicle control or M19 compound (20 μM, 24 hours). Differential expression analysis identified 633 upregulated and 543 downregulated genes following M19 treatment (Figure 3a). Gene set enrichment analysis (GSEA) of differentially expressed genes (DEGs) showed enrichment of pathways involved in apoptosis, the UPR, ROS signaling, and mTORC1 and p53 signaling (Supplementary Figure 3a). These pathways may underlie the toxicity that we observed in cultured tumor cells after M19 treatment and are consistent with pathways altered by inhibition of translation initiation complex components (14). Ingenuity pathway analysis (IPA) of these DEGs further identified activation of oxidative stress response pathway genes associated with NRF2 as well as UPR and WNT/beta-catenin signaling pathway genes (Supplementary Figure 3b, Supplementary Table 1). Quantitative PCR of 14 upregulated and 2 downregulated genes confirmed these expression changes (Supplementary Figure 3c). Among downregulated pathways were E2F target genes and genes involved in the G2M cell cycle control, changes confirmed by qPCR analysis (Supplementary Figure 3d). Treatment of melanoma cells with M19-6 phenocopied changes seen after M19 treatment, the most notable being in UPR (ATF4, DDIT3, and ATF3) and ROS (ERN1, HERPUD1, HHMOX, and c-Jun) signaling (Supplementary Figure 3e).

**Figure 3.**
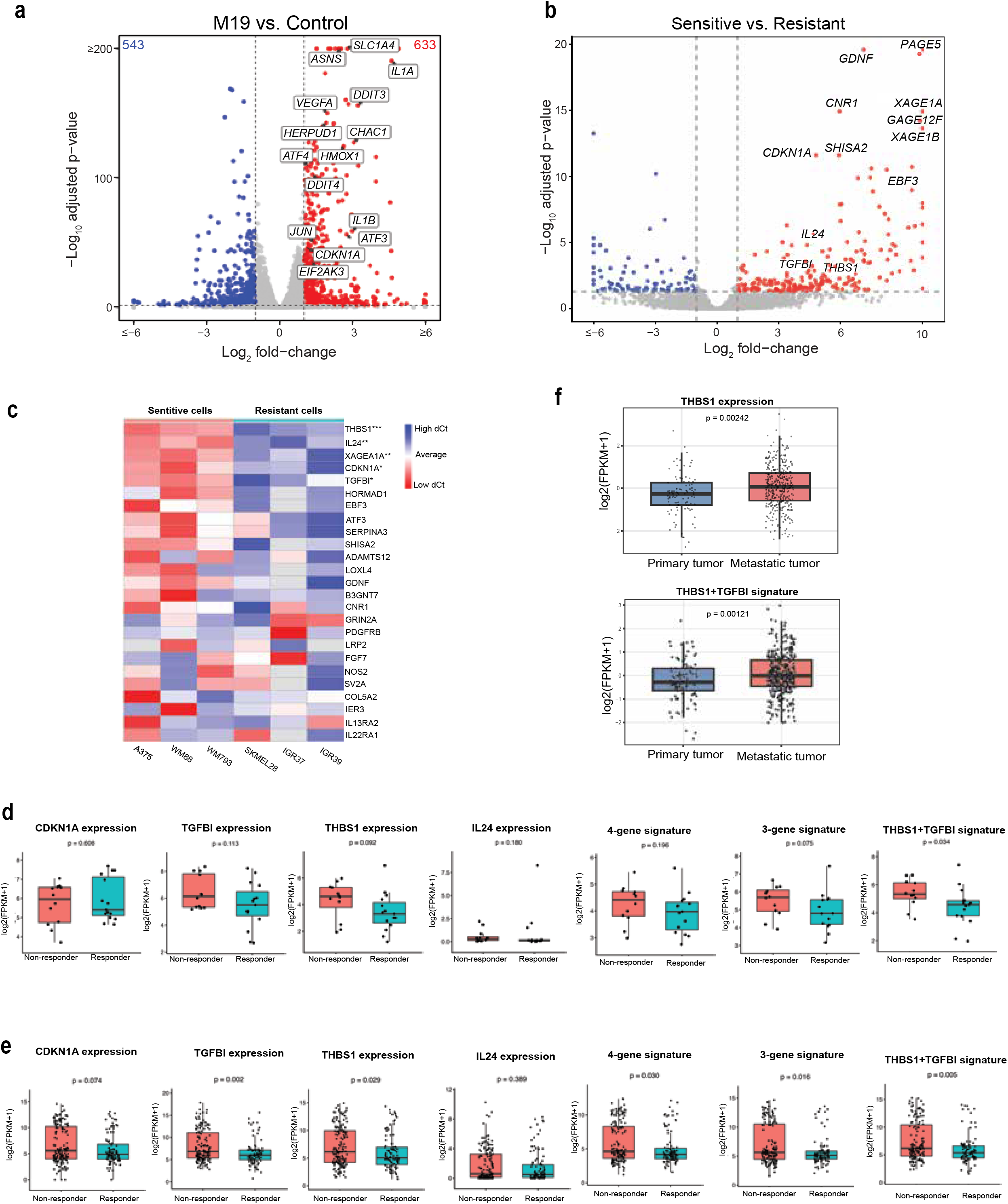
RNA-Seq analysis of M19-treated melanoma and characterization of M19 analogs. **(a)** Volcano plot comparing the transcriptomes of M19-treated and control A375 samples. The numbers of up- and down-regulated genes are indicated in the top corners. Representative genes that were up- or down-regulated upon treatment are denoted. **(b)** Volcano plot comparing the transcriptome from melanoma cell lines (CCLE). Differentially expressed genes between sensitive and resistant groups were analyzed using DESeq2. Top 20 ranked genes were labeled on the plot. **(c)** Heatmap showing qPCR of candidate biomarkers in M19-6–sensitive and resistant human melanoma cell lines. Gene expression was normalized to the reference gene and compared between sensitive and resistant groups. **(d)** Expression of THBS1, TGFβI, IL24, CDKN1A, and the indicated combined signatures (a 4-gene signature consisting of THBS1, TGFβI, IL24, and CDKN1A; a 3-gene signature consisting of THBS1, TGFβI, and CDKN1A; and a THBS1+TGFβI signature) was compared between responders and nonresponders in the GSE78220 cohort. **(e)** Expression of individual M19-6 markers and combined gene signatures was compared between responders and nonresponders across six pre-treatment melanoma anti–PD-1 cohorts. Patients were grouped by anti–PD-1 response status, and group differences were assessed using Wilcoxon test. Individual patient values are shown as black dots. **(f)** TCGA-SKCM RNA-Seq data were used to compare expression of THBS1 (upper panel) and a THBS1+TGFβI signature (lower panel) between primary and metastatic melanoma samples.

To determine whether sensitivity to M19-6 series may be linked with specific markers, we monitored changes in gene expression that distinguish responsive melanoma cell lines (A375, SKMEL5, WM88, WM793) and non-responsive melanoma cell lines (IGR37, IGR39, SKMEL28, MEWO). RNA-Seq data from CCLE (35) was used to compare gene expression between the sensitive and the resistance melanoma cell lines. Differential expression analysis identified 238 genes that were upregulated in sensitive melanoma cell lines and 93 genes that were upregulated in the melanoma resistant cell lines (Figure 3b). Of those, independent analyses confirmed that THBS1, IL24, CDKN1A, TGFβI and XAGE1A were upregulated in the melanoma cells that were sensitive to M19-6 treatment (Figure 3c).

To further explore the clinical relevance of these putative biomarkers, we analyzed the expression level of these genes in melanoma patients that were treated with anti-PD1 therapy. While THBS1 and TGFβI expression was elevated in non-responders, the expression of CDKN1A was elevated in the responders (GSE78220; (36) (Figure 3d). Extending the analysis to six additional melanoma patient cohorts that were treated with anti-PD1 therapy (36-38), revealed a similar trend, whereby expression of THBS1 and TGFβI was noted in non-responders (Figure 3e). These observation points to the possible use of these biomarkers for stratification of patients for M19-6 treatment, which could augment the effect of PD1 therapy.

Notably, TCGA-SKCM RNA-Seq data were further used to compare the gene expression between primary melanoma and metastatic melanoma samples. THBS1 and the combined “THBS1+TGFβI” signature was significantly higher in metastatic tumors than in primary tumors (Figure 3f). In contrast, IL24 and CDKN1A were significantly lower in metastatic tumors, while TGFβI alone was not (Supplementary Figure 4d). These findings suggest that the THBS1+ TGFβI signature may serve as markers for identifying metastatic melanoma patients that could benefit from M19-6 therapy.

### Translatome analysis of M19-6-treated cells

To investigate the impact of M19-6 on the transcriptome-wide collection of actively translated mRNA (translatome), we conducted ribosome profiling (Ribo-Seq) in A375 cells treated for 8 hours with M19-6 or vehicle control (Figure 4a). Ribo-Seq coupled with RNA-Seq enables transcriptome-wide assessment of translation efficiency of individual mRNAs (30). On average, we obtained approximately 37 million and 10 million mapped reads for Ribo-Seq and RNA-Seq, respectively, targeting coding regions (CDS) of the transcriptome. Micrococcal Nuclease (MNase) digestion generated ribosome footprints with a central peak around 33 nucleotides (nt) (Supplementary Figure 4a). As anticipated, metagene plots of Ribo-Seq data displayed robust accumulation of ribosome footprints within mRNA CDS regions (Supplementary Figures 4b, 4c). Principal component analysis (PCA) of Ribo-Seq and RNA-Seq libraries indicated reproducibility among independent replicates, with clear separation between M19-6-treated and control samples (Figure 4b). We detected 11,468 and 11,255 highly expressed mRNAs (RPKM > 1) in Ribo-Seq and RNA-Seq datasets, respectively, using the longest translon for each gene (Supplementary Table 2).

**Figure 4.**
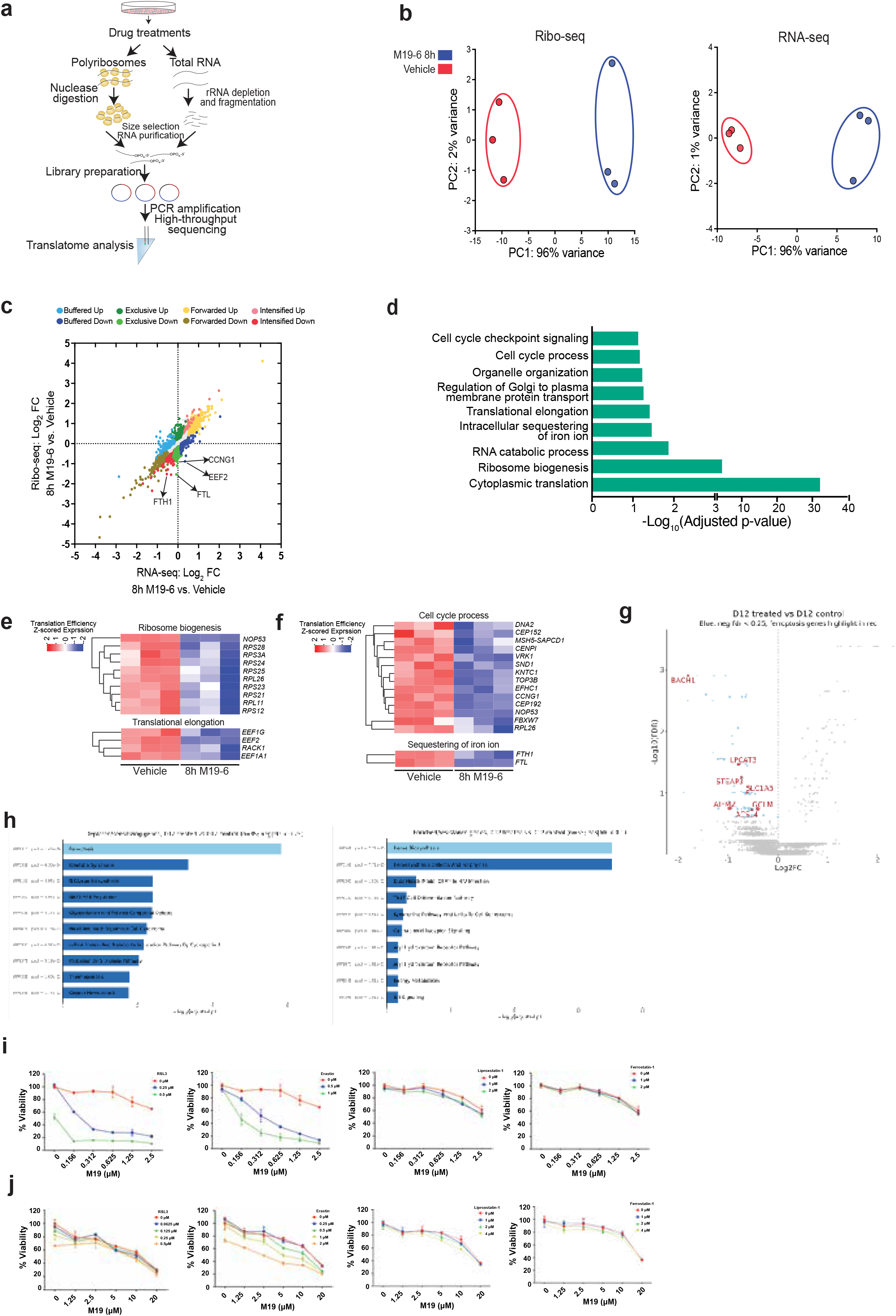
Translatome of M19-6 compound in A375 cells. **(a)** Schematic illustrating the experimental workflow for RNA-Seq and Ribo-Seq library preparation and sequencing in A375 cells. Biological replicates of A375 cells were treated with a control compound or M19-6 for eight hours prior to analysis. **(b)** Principal Component Analysis (PCA) of Ribo-Seq (left) and RNA-Seq (right) datasets, showing clear segregation by treatment. Each circle represents a biological replicate, with treatments color-coded for distinction. **(c)** Scatter plot depicting the relationship between fold changes (FC) in RNA-Seq and Ribo-Seq data for control versus M19-6-treated cells. Statistically significant mRNAs (p-adj < 0.05, Log2 FC > 0.2 or < -0.2) are categorized and color-coded as follows: *Intensified Up* (increased transcription and translation efficiency [TE]), *Intensified Down* (decreased transcription and TE), *Forwarded Up* (concurrent increases in mRNA and RPF with unchanged TE), *Forwarded Down* (concurrent decreases in mRNA and RPF with unchanged TE), *Exclusive Up* (increase in RPF and TE without mRNA change), *Exclusive Down* (decrease in RPF and TE without mRNA change), *Buffered Up* (decreased transcription/increased TE), and *Buffered Down* (increased transcription/decreased TE). **(d)** Gene Ontology (GO) enrichment analysis of translationally down-regulated mRNAs in M19-6-treated cells compared to control, with p-values adjusted for multiple comparisons via the Benjamini–Hochberg method. Down-regulated mRNAs with p-adj < 0.1 and -0.5 < Log2 FC < 0.5 were included in this analysis, with n = 49. **(e)** heat map of genes implicated in ribosome biogenesis which were enriched **(f)** heat map of genes implicated in cell cycle control which were enriched. **(g)** Volcano plot representation of the top candidate genes identified from the genome-wide CRISPR knockout screen in A375 melanoma cells treated with M19-6. **(h)** CRISPR knockout screen identifies genetic sensitizer (left)/resistance mediators (right) to M19-6 in melanoma cells. Enriched hits were subjected to pathway analysis. **(i)** A375 cells were treated with indicated concentrations of M19-6 and the ferroptosis inducer (RSL3, Erastin) or the inhibitor (Liproxstatin-1, ferrostatin-1) for 6 days before assessed for cell viability using CellTiter-Glo. **(j)** A375 spheroids were treated with indicated concentrations of M19-6 and the ferroptosis inducer (RSL3, Erastin) or the inhibitor (Liproxstatin-1, ferrostatin-1) for 6 days before assessed for cell viability using CellTiter-Glo 3D. The relative viability was normalized to vehicle-treated control.

To evaluate translation efficiency in control and M19-6-treated A375 cells, we applied the deltaTE (ΔTE) method (39), an enhanced version of DESeq2 optimized to assess translation efficiency from changes in Ribo-Seq and RNA-Seq data (39) (Figure 4c, Supplementary Table 3). Our analysis identified 104 mRNAs whose translation efficiency was down-regulated by M19-6 treatment (Figure 4c). Notably, most translation efficiency changes fell into the “Exclusive” category, in which mRNA levels remained unchanged, but Ribo-Seq reads were significantly up- or down-regulated (Figure 4c). These findings imply that M19-6 impacts gene expression primarily through translational regulation. mRNAs encoding proteins associated with ribosomal biogenesis, the cell cycle, and translational elongation were among gene ontology (GO) groups that were translationally down-regulated by M19-6 in A375 cells (Figure 4d, 4e, 4f; p-adj < 0.05 and log2 fold change in TE < -0.5). Changes in genes implicated in ferroptosis were also noted (Figure 4c and 4f). The finding that ferroptosis may be associated with M19-6 effectiveness led us to explore whether altering cells susceptibility to ferroptosis would enhance degree of cell death.

Whole-genome CRISPR knockout screen was performed to identify genes that may enhance the effectiveness of sensitivity or resistance of M19-6 on A375 melanoma cells. Genes involved in lipid peroxidation and redox homeostasis, including ACSL4, LPCAT3, and KEAP1, were identified as a key modulators of the M19-6 response (Figure 4g), suggesting that cellular susceptibility to oxidative stress plays an important role in determining treatment of sensitivity of melanoma cells to M19-6. Pathway analysis revealed ferroptosis as a prominently enriched pathway among genes whose loss increased sensitivity to M19-6 treatment (Figure 4h, left panel). In contrast, resistance to M19-6 revealed enrichment of heme biosynthesis pathway components (Figure 4h, right panel),

Addition of ferroptosis inhibitor (ferrostatin, liproxstatin) or inducer (Erastin, RSL3) to melanoma cells treated with M19-6 revealed enhanced cell death in the presence of the ferroptosis activator, but not inhibitor, in 2D culture (Figure 4i) and 3D spheroides (Figure 4j).

### M19-6 inhibits melanoma tumor formation and prevent melanoma metastasis

Assessment of M19-6 stability was conducted in mouse hepatocytes. We found that M19-6 exhibited a greater T1/2 (19.27 min) and a metabolic stability of 12.92% after 30 min, relative to 11.33 for Verapamil, a drug commonly used to treat hypertension (Supplementary Table 4).

To evaluate the effectiveness of M19-6 on melanoma growth *in vivo*, we used three melanoma models. The NRAS mutant melanoma selected for this analysis was SW1 cells (40,41) that were inoculated in C3H mice, the B16F10 melanoma represented MAPK amplification without upstream BRAF or NRAS mutation (42) and the YUMM1.5 melanoma which harbors BRAF mutation (43). Administration of M19-6 (90mg/kg daily injection) effectively decreased tumor growth in these three melanoma models (Figure 5a, 5b, 5c; Supplemental Figure 5a, 5b, 5c). Growth of YUMM1.5 melanoma tumors was also attenuated by the M19-6, (>50% Figure 5c; Supplemental Figure 5c), an expected outcome given the greater activity of M19-6 in NRAS-mutant melanomas. Notably, M19-6 treatment sufficed to inhibit spontaneous melanoma metastasis to the lungs, which was assessed in the SW1 model (Figure 5d).

**Figure 5.**
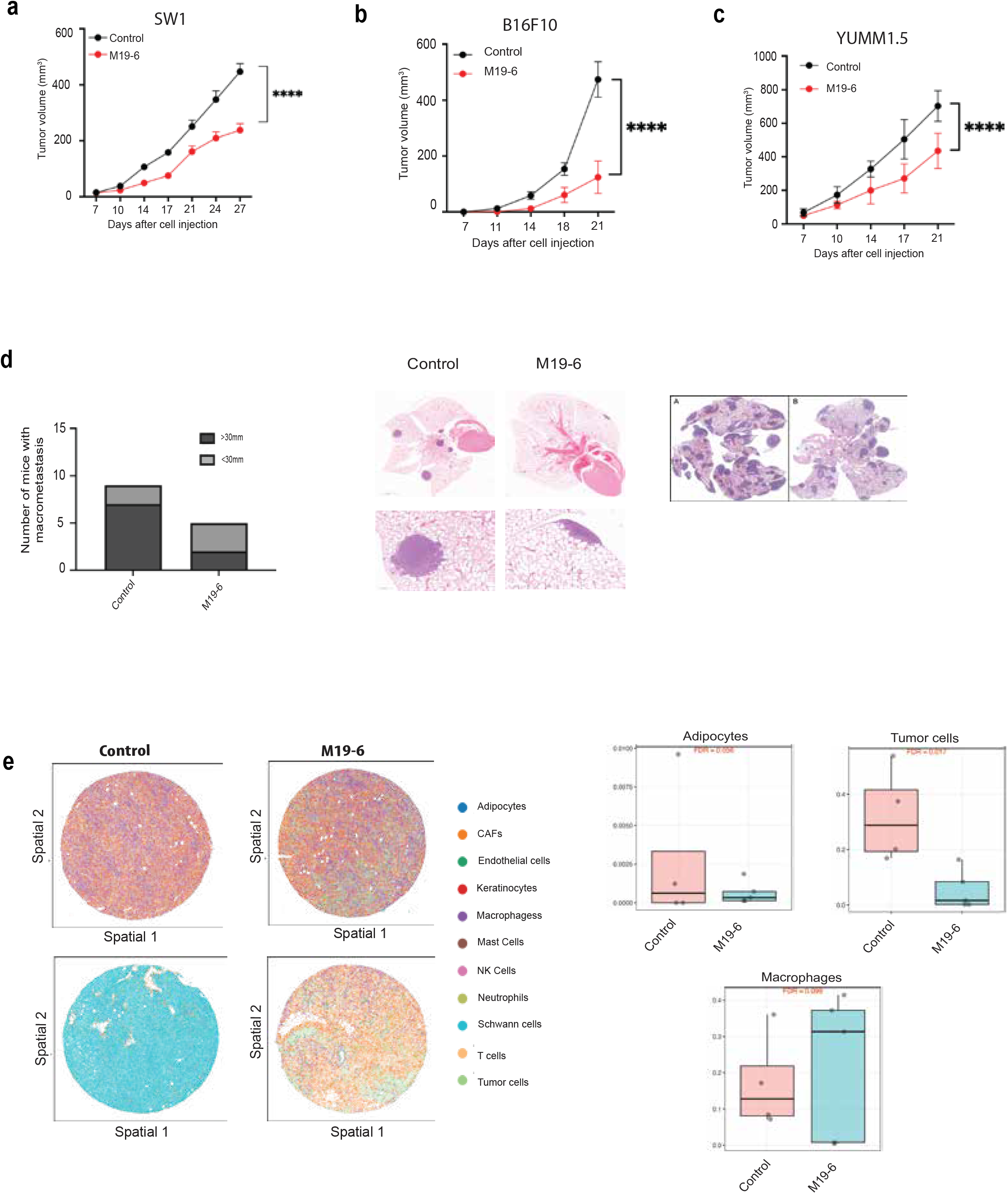
M19-6 inhibits melanoma growth and metastasis in vivo. **(a)** SW1 cells grafted into 8-week-old C3H/HeNHsd mice (n=10 per experimental group) subjected to treatment with M19-6. Shown is the growth of SW1 tumors. **(b)** B16F10 cells were inoculated in C57BL/6 mice (n=10 per experimental group) that were subjected to treatment with M19-6. Shown is the growth of B16F10 tumors. **(c)** YUMM1.5 cells were inoculated in C57BL/6 mice (n=10 per experimental group) for 7-10 days before emerging tumors were subjected to treatment with M19-6. Size of the tumors at indicated time points is shown. **(d)** SW1 cells were inoculated s.c into 1-year-old C3H/HeNHsd mice (n=10 per experimental group), and a week later, when tumors were palpable, M19-6 was administered for the indicated time periods. Lung metastasis was quantified (number and size) and H&E images were taken. **(e)** Spatial distribution of immune cell types in tumor samples and their quantification (Control, n=3; M19-6, n=3). Data were analyzed by unpaired t-test. *P < 0.05, **P < 0.005, ***P < 0.001, ****P < 0.0001 by two-tailed t-test or two-way ANOVA.

A recent study suggested that treatment with SBI-640756 (the parental M19-6 drug) induces anti-tumor immune phenotypes (26). We thus performed spatial analyses of tumors to identify changes in immune cell infiltration following M19-6 treatment. Notably, we identified difference in the composition of the immune cells in different areas of tumor (two different areas are shown per treatment; Control and M19-6) (Figure 5e, left panel). Notably, this analysis revealed increased infiltration of macrophages and reduced proportion of adipocytes within B16F10 tumors of mice treated with M19-6, compared with non-treated mice (Figure 5e, right panel). Immunophenotyping that was performed on YUMM 1.5 tumors, collected 12 days after tumor cell inoculation in C57BL/6J mice, confirmed increased abundance of macrophages within the tumors (Supplemental Figure 5d). These data suggest that administration of M19-6 increases abundance of macrophages within melanoma tumors, that may account for enhanced anti-tumor immunity.

## Discussion

Targeting the eIF4F complex in neoplasia presents challenges since eIF4F is essential for the viability of all cells. However, this apparent disadvantage may be exploited, as complex components are generally overexpressed in tumors relative to normal cells (7-9). Strong genetic evidence supporting the targeting of eIF4F has come from analysis of mice heterozygous for eIF4E, which are reportedly resistant to cancer development despite exposure to carcinogens (44). Among the small molecule inhibitors designed to target this complex, a limited number have advanced to clinical trials, with the eIF4A inhibitor eFT226 being a notable example (45). Thus, this study was designed to fill an unmet clinical need, leveraging our earlier analyses of SBI-756, a small molecule that interacts with eIF4G1 and dissociates the eIF4F complex (5).

Herein, we first discovered that a small molecule inhibitor, SBI-756, binds the MA3 domain of eIF4G1. This finding guided structure-based *in silico* screen that identified M19 and its analog M19-6, novel small molecule inhibitors of eIF4G1. Notably, compared to SBI-756, these compounds demonstrate dose-dependent suppression of melanoma cell proliferation in 2D and 3D culture models. Furthermore, they effectively disrupted the eIF4F complex and inhibited the interaction between eIF4G1 and eIF4A, underscoring their potential therapeutic efficacy. RNAs-eq analysis of melanoma cells treated with M19 revealed ROS, UPR, and apoptosis signaling activation, indicating that M19 induces cell death programs. Further assessment using Ribo-Seq approach led to the identification of ribosomal structure components that can be linked to growth arrest phenotypes, which were inhibited following treatment with the M19-6 compound, consistent with the sensitization of melanoma cells to cell death in culture, resembling the requirement of other eIF4F inhibitors for structural ribosomal proteins (22). Notably, this analysis also identified the ferroptosis protein FTH1 and FTL, that were confirmed to underlie M19-6 activity. Along these lines, unbiased search for pathways that may complement M19-6 activity, using whole-genome CRISPR screen, led to the identification of genes implicated in ferroptosis as top candidates.

Among the unmet clinical needs in clinical management of melanoma is the emerging resistance of tumors to therapy. Notably, M19-6’s ability to attenuate eIF4F complex assembly enables it to overcome therapy resistance, consistent with findings made with the SBI-756 (5). M19-6 was shown to be effective against BRAF- and MEK-resistant melanoma, confirming the need for complex dissociation to counteract therapy resistance.

Targeting of the eIF4F complex has been primarily achieved using small molecules that impede the activity of eIF4E or eIF4A subunits (4,9). Moreover, reluctance to target eIF4G1 has been justified by a lack of structural data or based on its “adaptor-like” function (46). Indeed, our earlier generation small molecule, SBI-756, is an effective inhibitor of the eIF4F complex, although its toxicity in culture and biophysical properties precluded its further development. Of the >140 SBI-756 analogs previously developed, none exhibited activity superior to SBI-756, while its pharmacokinetics were suboptimal. We have now overcome these limitations by identifying the eIF4G1 MA3 domain as the SBI-756 target and used structure-based analysis to identify and characterize a new generation of small molecule eIF4G1 inhibitors. Pharmacological inhibition of eIF4G1 with these inhibitors was more effective than genetic inactivation of the MA3 domain, likely due to the presence of the eIF4G2 isoform, which also harbors the MA3 domain (27). We anticipate that targeting the MA3 domain by M19-6 will impact both eIF4G isoforms, with limited toxicity, especially at low doses used in culture and the doses used in preclinical models.

Previous studies demonstrated that eIF4G1 inhibition perturbed the cell cycle and DNA repair processes (14), changes also observed in the presence of SBI-756 (5) and the M19 compound series. Disruption in cell cycle is a long-sought goal in cancer therapy, an endpoint that has guided the development of numerous inhibitors that have reached clinical evaluation; nonetheless, common to eIF4F inhibitors studied so far is that all must be used in combination with drugs that target other cellular pathways to achieve effective and sustained clinical outcomes, as exemplified by CDK4/6 inhibitors (47). Our discovery of M19-6 compounds highlights the opportunity for combination therapy, where the second drug could be a ferroptosis activator, although currently largely unavailable as effective in vivo drugs, their active development is expected to reach clinical evaluation in the near future.

In all, our work describes the development of the new generation of eIF4G1 inhibitors that target the MA3 domain of the eIF4G1 gene. Compared with B-RAF mutant, M19-6 is more effective in NRAS mutant melanoma and overcomes resistance to targeted therapies as well as limits metastasis – a prevalent unmet clinical needs in melanoma management. Exemplified by M19-6, this new class of inhibitors offers advantages over existing eIF4F inhibitors and merits further evaluation in preclinical and clinical models.

## Materials and Methods

### Mouse tumor models and ethics

All animal studies were approved by the Institutional Animal Care and Use Committee (IACUC # 21-064) of Sanford Burnham Prebys Medical Discovery Institute and Cedars-Sinai Medical Center (Protocol: IPROTO202400000350) and complied with ethical animal testing and regulations. Six-week-old and 1-year-old male C3H/HeNHsd mice were purchased from Inotiv and six-week-old male C57BL/6J mice were purchased from Jackson Laboratories, allowed to acclimatize for one week. SW1 cells (8×10^5^), B16F10 cells (2.5 × 10^5^), YUMM1.5 cells (1 × 10^6^), suspended in 200 μL sterile PBS were injected subcutaneously into the flanks of mice. When tumor sizes reached ∼150 mm^3^, mice were sorted into different groups and treated with IP injection of Vehicle or M19-6 (90 mg/kg, once per day). Body weight and tumor volume were measured twice a week. Tumor size was measured with linear calipers and calculated using the formula: ([length in millimeters × (width in millimeters)^2^]/2). Mice were sacrificed after 4 weeks, and tumors and lungs were fixed in Z-Fix and paraffin-embedded for immunohistochemistry analysis.

### Cell culture

All human and mouse melanoma cells were maintained in high-glucose Dulbecco’s modified Eagle’s medium (DMEM; Cytiva, Marlborough, MA) with 10% fetal bovine serum (FBS) and 1% penicillin-streptomycin (Cytiva). Cell cultures were maintained at 37°C in 5% CO_2_.

### Reagents

Compounds used in this study were purchased from ChemBridge (San Diego, CA), Enamine (Monmouth Junction, NJ), and MolPort (Beacon, MA), as indicated. Chloroquine was obtained from Sigma-Aldrich. Compounds were dissolved in DMSO and stored as a 10 mM stock solution at -20°C. Each compound was freshly prepared from the stock solution before use. Compounds used in animal studies were synthesized by Pharmaron (Qingdao, China) at >96% purity.

### Cell viability assay

All cell lines were seeded in 96-well white plates (5,000 cells in 100 μl per well) or 384-well white plates (500 cells in 50 μL per well) and allowed to attach overnight. Test compounds were serially diluted from 10 mM stock solutions and added to cells. Assays were performed in triplicate, and each microplate included media and DMSO control wells. Cell viability was assessed 72 h after treatment using CellTiter-Glo (Promega) according to the manufacturer’s protocol. For the spheroid growth inhibition assay, cells were seeded in 96-well (1000 cells in 100 μl per well) or 384-well (250 cells in 50 μL per well) white Corning spheroid microplates and allowed to grow 24 h to form spheroids. Cell viability was assessed 5 days later using CellTiter-Glo 3D (Promega) according to the manufacturer’s protocol. In both cases, cell growth inhibition was calculated as a percentage of DMSO-treated controls.

### Colony formation assay

Cells were plated in triplicate (500 cells/well) in 6-well plates and allowed to grow overnight before compounds were added. After 1–2 weeks, depending on the cell line, colonies were stained with Crystal Violet solution (Sigma-Aldrich) for 30 min. Plates were rinsed with water, and images were acquired by scanning. Colony number and size were analyzed using Image J software. Colony formation efficiency was calculated relative to the number of colonies in control (DMSO)-treated wells.

### Antibodies, Immunoprecipitation, and Western blot analyses

Cells were rinsed with PBS and lysed in lysis buffer (50 mM Tris-HCl, pH 7.4, 150 mM NaCl, 1% NP-40, 1 mM EDTA) containing protease inhibitors and a phosphatase inhibitor cocktail (Thermo Scientific, Waltham, MA). Protein concentration was determined using Coomassie Plus Protein Assay Reagent (Thermo Scientific). Immunoprecipitation (IP) of eIF4A was performed from cell lysates (300 μg) incubated overnight with 10 μg of anti-eIF4A antibodies (BioLegend, San Diego, CA, cat# 666202), followed by incubation with Protein A/G PLUS-Agarose beads (Santa Cruz, SC-2003) for 1 h. Beads were then washed 4 times with lysis buffer, and IP’d proteins were eluted with SDS loading buffer and analyzed by Western blot. Equal amounts of cell lysate proteins (20-50 μg) or proteins eluted from agarose beads were separated on SDS-PAGE and transferred to polyvinylidene difluoride (PVDF) membranes (PerkinElmer Life Sciences). Antibodies against eIF4G1, eIF4E, and 4EBP1 were purchased from Cell Signaling Technology (cat# 8701, cat# 2067, cat# 9644). Actin and HSP90 antibodies were obtained from Santa Cruz Biotechnology (cat# sc-58673, cat# sc-13119). Secondary antibodies were goat anti-rabbit Alexa Fluor 680 (cat# A21076), goat anti-mouse Alexa Fluor 680 (cat# A21057), and goat anti-rat Alexa Fluor 680 (cat# A21096; Invitrogen).

### Construction of EIF4G1 tiling sgRNA library

The custom EIF4G tiling sgRNA library consisting of 947 unique sgRNAs was designed using the Broad Institute Genetic Perturbation Platform (GPP) sgRNA designer tool (48,49). Library sequences were synthesized on an array platform (Genscript, NJ) and then cloned into the lentiGuide Puro vector (Addgene: #52963) using Golden Gate Assembly. The library plasmid was amplified using ElectroMAX™ Stbl4™ Competent Cells (Invitrogen, cat # 11635018). To ensure representation, the number of bacterial colonies was maintained at >500 times the number of sgRNA sequences in the library. sgRNA sequences in the library are deposited as indicated in the data availability section.

### Selection of SBI-756-resistant clones and identification of corresponding gRNAs

The pooled tiling CRISPR library virus was produced using HEK293T cells transfected with pMD2.G and psPAX2 in the presence of Polyethylimine (PEI). A375 cells (2 × 10^6^) were then transduced with the pooled library lentivirus in DMEM complete medium and 0.8 μg/ml polybrene. Twenty-four hours later, virus, polybrene, and medium were removed, and cells were plated in fresh culture medium. Forty-eight hours after that, puromycin was added at 1⍰μg/ml to select for sgRNA library-expressing cells. The multiplicity of infection (MOI) was adjusted to 0.3-0.4 to prevent >1 sgRNA infection per cell. Cells were cultured at >1,000X of the library size to ensure adequate sgRNA representation. Cells were treated with indicated SBI-756 concentrations for 2-4 weeks until resistant clones emerged. Genomic DNA extraction from drug-resistant cells was performed using the Zymo QuickDNA™ Midiprep Plus Kit (Zymo Research, Cat # D4068).

### gRNA sequencing data processing

A two-step PCR-amplification for sequencing library preparation was conducted with TaKaRa Ex Taq™ Polymerase (TaKara, Cat. RR001) and custom primers, to achieve adequate sequencing coverage in 1 × 75bp single-end reads, following published guidelines (50). Barcoded libraries were pooled and sequenced using an Illumina Hiseq 500 platform.

### CRISPR screen data analysis

The sequence and Pfam domains of the longest transcript of EIF4G1 gene (ENST00000424196.5) were obtained from the Ensembl database. Raw reads were sequentially trimmed for 3’ and 5’ adapters using Cutadapt v2.8 with parameters “-j 4 -e 0.15 -a GTTTTAGAGCTAGAAATAGCAAGTTAAAATAAGGCTAGTCCGTTATCAAC” and “-j 4 -e 0.15 -m 18 –g TTTATATATCTTGTGGAAAGGACGAAACACCG” respectively. The trimmed reads were aligned to EIF4G1 longest transcript sequence using STAR v2.7.0d_0221(51)) with parameters “--runThreadN 8 --alignEndsType EndToEnd --outFilterMismatchNmax 0 -- outFilterMultimapScoreRange 0 --outReadsUnmapped Fastx --outSAMtype BAM Unsorted-- outFilterMultimapNmax 1 --outSAMunmapped None --outFilterScoreMinOverLread 0 -- outFilterMatchNminOverLread 0 --outFilterMatchNmin 20 --alignSJDBoverhangMin 1000 -- alignIntronMax 1”. Signal bigwig files were generated using Deeptools bamCoverage v3.2.0 (52) with parameters “—normalize using CPM --exactScaling”.

### Selection of single-cell gRNA clones

Individual gRNA235 or gRNA622 virus was produced using 293T cells following standard protocols and used to infect A375 cells following puromycin selection (1lμg/mL). Single-cell clones were generated by serial dilution (0.2 cell/100 μL) in 96-well plates. The resulting single-cell clones were treated with SBI-756, and resistant clones were selected. Genomic DNA was extracted from selected clones, and the region corresponding to the gRNA was PCR-amplified and sequenced to confirm the specific deletion at the targeted region.

### m^7^GTP pull-down assay

Cells growing in 100 mm plates and treated with indicated compounds were washed with cold PBS, collected, and lysed in lysis buffer [50 mM MOPS/KOH (pH 7.4), 100 mM NaCl, 50 mM NaF, 2 mM EDTA, 2 mM EGTA, 1% NP40, 1% Na-DOC, 7 mM β-mercaptoethanol containing protease inhibitors and a phosphatase inhibitor cocktail]. Lysates were incubated for 20 min with m^7^-GDP-agarose beads (Jena Bioscience, Jena, Germany) with rotation and washed three times with wash buffer (50 mM MOPS/KOH, pH 7.4, 100 mM NaCl, 50 mM NaF, 0.5 mM EDTA, 0.5 mM EGTA, 7 mM β-mercaptoethanol, 0.5 mM PMSF, 1 mM Na3VO4 and 0.1 mM GTP). Bound proteins were eluted by incubating beads in a loading buffer. Materials pulled down by m^7^-GDP-agarose were analyzed by Western blot.

### Monitoring of newly synthesized proteins by puromycin labeling

For labeling experiments, A375 cells were seeded in 6-well plates (3 × 10^5^ cells per well). After 24 h, cells were pre-incubated for 1 h with different compounds prior to treatment with 3 μg/mL puromycin for 30 min. Cells were washed 2 times with ice-cold PBS and lysed with lysis buffer. Cell lysates were subjected to Western blot analysis.

### Histology and immunofluorescence

Tumors and lung were collected by the end of the experiment. Tissues were fixed with 4% formalin, washed with PBS, embedded in paraffin and cut into 5 μM-thick sections and stained with H&E.

### Tumor digestion

Tumors were harvested at day 12 after tumor injection and digested with 100 μg/mL DNase I (sigma) and 1mL of collagenase D (Roche) at 37°C for 1 hour. To generate single-cell suspension, the tumor digested was passed through a 70 μm cell strainer. The cells were washed with PBS and incubated with the indicated antibodies for flow cytometry.

### Flow Cytometry analysis of tumor cells

Tumor single cells were washed with FACS staining buffer and incubated with antibodies (dilution 1:200) at 4°C for 20 min and fixed using 1% of formaldehyde in PBS for 20 min, followed by washing before resuspending in FACS staining. The antibodies used were: NK1.1 (Clone: OK136), CD8 (Clone: 53-6.7), CD4 (Clone: RM4-5), Lag3 (Clone: C9B7W), CD69 (Clone: H1.2F3), B220 (Clone: RA3-6B2), CD44 (Clone: IM7), CD45 (Clone: 104), CD279 (Clone: 29F.1A12), MHC II (Clone: M5/114.15.2), CD11b (Clone: M1/70), PDCA-1 (Clone: 129c1), CD11c (Clone: N418), MHC I (Clone: AF6-88.5), F4/80 (Clone: BM8), CD206 (Clone: C068C2), GR1 (Clone: RB6-8C5), CD80 (Clone: 16-10A1), TCF-1 (Clone : S33-966), TIM-3 (Clone : B8_2C12). All the data were collected on an BD FACSymphony A5 and analyzed using Flowjo Software.

### Analysis of the cell cycle

A375 cells that express the gRNA constructs were subjected to FACS analysis following their trypsinization, fixation (60 min in 70% ice-cold ethanol), and washes in phosphate-buffered saline, followed by incubation for 30 min at 25°C in propidium iodide (PI) buffer (10 mM Tris-HCl, pH 7.4, 5 mM MgCl2, 50 μg of PI per ml, and 10 μg of RNase A/ml). Stained cells were acquired using the FACSort flow cytometer (BD Biosciences), and DNA content was analyzed using FlowJo software.

### Analysis of apoptosis and ferroptosis

A375 cells were seeded in 96- or 384-well white plates, treated with different concentrations of indicated compounds, and apoptosis was measured at different time points using Caspase-Glo 3/7 (Promega, G8091), according to the manufacturer’s protocol. Caspase-Glo 3/7 3D (Promega, G8981) was also used to analyze A375 spheroid apoptosis induced by different treatments. A375 cells were treated with M19-6 alone or in combination with a ferroptosis inducer or inhibitor. After treatment, cell viability was quantified using CellTiter-Glo, and relative viability was normalized to vehicle-treated controls.

### Reverse transcription and real-time qPCR

Total RNA from DMSO- and compound-treated cells and tumors was extracted using a total RNA miniprep kit (Zymo Research). cDNA was synthesized using a High-Capacity cDNA Reverse Transcription Kit from Life Technologies, according to the manufacturer’s protocol. Real-time PCR was performed on a Bio-Rad CFX Connect Real-Time System using SsoAdvanced Universal SYBR®Green Supermix from Bio-Rad. H3A served as an internal control. Quantitative PCR reactions were performed in triplicate. PCR primers were designed using PrimerBank (http://pga.mgh.harvard.edu/primerbank) and are listed (Supplementary Table 3).

### Polysome profiling

A375-gRNA235 and A375-gRNA622 cells were seeded in 150 mm plates and treated with 0.2 μM SBI-756. After 24 h, cells were harvested and lysed in hypotonic lysis buffer (5 mM Tris HCl, pH 7.5, 2.5 mM MgCl2, 1.5 mM KCl, 100 μg/ml cycloheximide, 2 mM DTT, 0.5% Triton, 0.5% sodium deoxycholate). Polysome-profiling was carried out as described (31). Fractions were collected, and RNA was extracted using TRIzol, based on the manufacturer’s instructions. Experiments were done in independent triplicate samples.

### RNA-Seq

Total RNA was extracted using a total RNA miniprep kit (Sigma) with the supplied On-column DNase I for the digestion step. Libraries were prepared from isolated total RNA using the QuantSeq 3’ mRNA-Seq Library Prep Kit FWD for Illumina (Lexogen, Vienna, Austria). Barcoded libraries were pooled and single end-sequenced (1 × 75) on the Illumina NextSeq 500 system, using the High output V2.5 kit (Illumina Inc., San Diego, CA). Read data was processed and multiplexed with the BlueBee Genomics Platform (BlueBee, San Mateo, CA).

### RNA-Seq data processing

The Illumina adapter, polyA/polyT and low-quality sequences were trimmed from raw RNA-seq reads using Cutadapt v2.3. The trimmed reads were then aligned to human genome version hg38 and Ensembl gene annotations version 84 using STAR v2.7.0d_0221(70) with parameters adopted from ENCODE long RNA-seq pipeline (https://github.com/ENCODE-DCC/long-rna-seq-pipeline). Gene-level counts and normalized expression (TPM) were obtained using RSEM v1.3.1 (3). Trimmed reads and alignment qualities were assessed using FastQC v0.11.5 (https://www.bioinformatics.babraham.ac.uk/projects/fastqc/) and MultiQC v1.8 (53). Genes with RSEM-estimated counts equal to or greater than 5 times the total number of samples were retained for downstream analysis. Differential expression comparison was performed using DESeq2 v1.22.2. Gene set enrichment analysis was performed using the GSEA app v4.3.2 (54)

### Ribosome Profiling

Cultured cells were lysed in-dish using 250 μL of 2X lysis buffer, containing 40 mM Tris-HCl (pH 7.4), 300 mM NaCl, 10 mM MgCl_2_, 200 μg/mL cycloheximide (CHX), and 2 mM dithiothreitol (DTT), supplemented with Complete™ Protease Inhibitor Cocktail. Lysed cells were scraped into an Eppendorf tube (∼500μL), 5 μL of RiboLock RNase inhibitor (ThermoFisher Scientific, Cat#EO0382), and 5 μL of Turbo DNase (ThermoFisher Scientific, Cat#AM2238) were added to each lysate. The mixture was vortexed briefly (5 sec), incubated on ice for 10 minutes, and then centrifuged at 16,000× g for 10 minutes at 4°C. The optical density was recorded at A260 using a Nanodrop to assess concentration. Samples were stored at -80°C, with a portion reserved for RNA-Seq prior to Micrococcal Nuclease (MNase) treatment.

For ribosome profiling, lysates underwent MNase digestion (25U MNase (Sigma) per A260 unit) for 45 minutes at room temperature. The digestion was halted by adding EGTA. Monosomes were isolated by layering the digested polysomes over a 10-50% sucrose gradient, followed by centrifugation at 36,000 rpm for 3 hours at 4°C (Beckman Coulter). RNA was then extracted from the monosomes fraction using Trizol (Invitrogen) as per the manufacturer’s guidelines. In parallel, total RNA was extracted with Trizol from a portion of the untreated lysate for RNA-seq analysis.

For Ribo-seq library preparation, RNA samples were size-selected on a 15% TBU-Urea gel, with fragments between 25–40 nt isolated. The gel slices containing these fragments were incubated in 700 μL of RNA gel extraction buffer (300 mM NaOAc pH 5.5, 1 mM EDTA, 0.25% SDS) on dry ice, then rotated overnight at room temperature. Footprint fragments were recovered via isopropanol precipitation. The ribosome footprints were dephosphorylated with T4 Polynucleotide Kinase (T4 PNK) for 1 hour at 37°C in 1X T4 PNK buffer, with the addition of 10 U SUPERase*In and 5U T4 PNK (10 U/μL). Ligation was achieved by adding the ligation mix (17.5% PEG-8000, 1X T4 RNA ligase buffer, 100 U T4 Rnl2(tr) K227Q (200 U/μL), and 1 μM pre-adenylated linkers) to the samples. Ligated RNA was separated on a 10% TBE-Urea gel, extracted, and purified using RNA extraction buffer and isopropanol. Reverse transcription was conducted with SuperScript III (Invitrogen), and samples were circularized with CircLigase I (Lucigen). A round of rDNA depletion was performed using biotin-labeled antisense probes and Dynabeads MyOne Streptavidin C1 (55). The final libraries were size verified on an 8% TBE gel, quantified, and sequenced on an Illumina Platform.

For RNA-seq library preparation, the same protocol was applied, with modifications. The NEBNext® rRNA Depletion Kit (Human/Mouse/Rat) was used for rRNA removal, and the rRNA-depleted RNA was fragmented using the NEBNext® Magnesium RNA Fragmentation Module. To process sequencing data, initial trimming of adapters from reads was conducted with Cutadapt 3.4, allowing no mismatches in adapter sequences, and discarding reads shorter than six nucleotides. Trimmed reads were then demultiplexed by sample-specific barcodes anchored at the 5’ end, with no mismatches permitted. Sequential alignment was performed against the human transcriptome, beginning with cytoplasmic and mitochondrial rRNAs and tRNAs, using Bowtie 1.3.0 (56). Reads with low quality or shorter than 25 nucleotides were removed. Remaining reads were aligned to the comprehensive transcriptome (Gencode v32(57) with Bowtie 1.3.0, allowing two mismatches. The SAM files generated were converted to BAM files, sorted, indexed, and processed to remove PCR duplicates with UMI-Tools 1.0.1 (58). The final BAM files were converted into an SQLite format for analysis in Trips-Viz, a dedicated platform for Ribo-seq data (59). For metagene profile generation, annotated start and stop codons of coding sequences were aligned across all principal transcript isoforms by read length. To estimate the A-site position of each read, we calculated the nucleotide distance from the CDS start to the most prominent peak upstream, adding three nucleotides to this value.

### Differential Expression Analysis

Count matrices for RNA-Seq and Ribo-Seq data were generated using the “Read Table” function in Trips-Vis (59), which quantifies reads that uniquely align to the CDS region of mRNAs based on inferred A-sites. A Ribo-Seq read contributes to the CDS count if its inferred A-site falls within the annotated start and stop positions of the CDS in a given transcript. The genomic coordinates for each read were assessed to ensure unambiguous mapping. Differential analysis was conducted with the deltaTE (ΔTE) package in R (version 1.34.0; (55), which extends DESeq2 by applying an interaction model. For each gene, adjusted p-values (p-adj) were computed to assess fold changes between two experimental conditions across Ribo-Seq, RNA-Seq, and TE levels, with a significance threshold of p-adj < 0.05 for each fold change. These fold changes allowed categorization of genes into eight groups, as described in the text. In the scatter plots for fold changes between Ribo-Seq and RNA-Seq (Figure 4C), genes with “NA” values in any of the comparisons (Ribo-Seq or RNA-Seq) were omitted. PCA plot values in Figure XB were also obtained using DESeq2.

### In silico screen for MA3-bound small molecules

To map SBI-756 binding to the MA3 structure, a blind docking technique (molecular docking without a defined target site) was first performed using Glide (60) and FRED docking software(80). This analysis revealed a potential site where SBI-756 formed stable bonds during a 30ns molecular dynamics simulation performed with the Desmond suite from Schrodinger (33).

To identify small molecule inhibitors of eIF4GI, we carried out a virtual screen of the MA3 binding pocket. Based on molecular docking of a large library with subsequent filtering (34), a screen with a library of 10 million drug-like compounds (deposited in the ZINC20 database; (61) was carried out. Compounds were docked in the pocket based on three docking software systems, namely, GlideSP (60), FRED (62), and Autodock (63), which were shown to improve hit rates (85, 86). Binding poses predicted by these systems were compared, and compounds with consensus predictions (binding poses within 3Å distances) were selected. These compounds were ranked based on Glide docking scores. A total of 54,500 compounds received a more favorable docking score than the predicted SBI-756 with the docking score of -4.7. As SBI-756 presents an active scaffold targeting the MA3 pocket, we developed pharmacophore models contained three hydrophobic features corresponding to each of the three separate SBI-756 ring systems—one hydrogen donor and one hydrogen acceptor corresponding to the nitrogen and oxygen atom of the quinolone group. These features were used to select 500 chemically diverse compounds. Next we applied Andrew’s scoring metric, which assesses how well a molecule binds to an average receptor, by assigning partial scores to each functional group in the molecule (64). The remaining compounds were ran through the Swiss-ADME platform to predict physicochemical properties and potential off-target effects (64). Compounds predicted to inhibit 5 or 6 CYP family enzymes, which are critical for drug metabolism and clearance, were filtered out. Compounds receiving predicted log P values (lipophilicity) <5 were selected, as highly lipophilic compounds tend to be more promiscuous and are minimally soluble (65).

Compounds with low solubility in aqueous media (log S < -6) were excluded to avoid low bioavailability. The remaining 150 compounds were clustered, and cluster representatives were selected. Clustering procedures ensure that the purchased compound set is as chemically diverse as possible, maximizing the likelihood of finding an active scaffold. 67 potential inhibitors that bind the MA3 pocket were selected for biological evaluation.

### Hepatocyte stability assay

A hepatocyte stability assay was performed by Pharmaron (Beijing, China). Briefly, mouse hepatocytes (0.5×10^6^/ml) were dispensed into microtubes and 10 μL aliquots incubated (10 min) with the test compound (100 μM) or Verapamil (positive control; 100 μM), followed by incubation with 6 volumes of cold acetonitrile with IS (100 nM alprazolam, 200 nM labetalol, 200 nM caffeine and 200 nM diclofenac) to terminate the reaction. Samples were then centrifuged (45 minutes at 3,220 g), and 100 μL aliquots of supernatants were diluted with 100 μL ultra-pure H_2_O for LC/MS/MS analysis.

### Genome-wide CRISPR screen to identify genes whose loss sensitizes cells to M19-6

A375 cells expressing Cas9 were transduced with the Brunello CRISPR knockout library at MOI < 0.3 and selected with puromycin to establish the baseline population. Cells were treated with 0.5 μM M19-6 or DMSO for 9 or 12 days. Genomic DNA was extracted, sgRNAs were PCR-amplified, and analyzed by next-generation sequencing. sgRNA counts were normalized and compared between treated and control groups to identify genes whose loss sensitized cells to M19-6. Sequencing data from the CRISPR screen were analyzed with MAGeCK v0.5.9.5 (66) using the Broad Brunello sgRNA library. Lane-level FASTQ files were grouped by sample, sgRNA counts were generated with automatic 5’ trimming, and gene-level testing was performed with control sgRNAs targeting non-essential genes for normalization (67). Genes depleted in treated cells were interpreted as candidate sensitizing hits, whereas genes enriched in treated cells were interpreted as candidate resistance hits.

For pathway analysis, depleted genes were defined using FDR < 0.25 and enriched genes using FDR < 0.1. Each gene set was analyzed separately with Enrichr (68) against the `WikiPathways_2024_Human` collection. Gene-level heatmaps were generated from normalized sgRNA counts by calculating treatment-versus-control log2 fold changes for each guide and aggregating guide effects to genes, with median aggregation used as the primary representation.

### Analyses of melanoma patients dataset

To evaluate the clinical relevance of M19-6-associated biomarkers, GSE78220 (anti–PD-1 dataset; (36) was analyzed for gene expression and treatment response. Based on anti–PD-1 response patients were grouped as responders or non-responders. Expression of candidate genes, including THBS1, TGFβI, IL24, and CDKN1A, was assessed in these patients. The expressions of THBS1, TGFβI, IL24, and CDKN1A were analyzed either individually or as combined signatures. The average expression of the following combinations was determined: a 4-gene signature consisting of THBS1, TGFβI, IL24, and CDKN1A; a 3-gene signature consisting of THBS1, TGFβI, and CDKN1A; and a THBS1+TGFβI signature. Individual gene expression values and signature scores were compared between responder and non-responder groups. Group differences were assessed using Wilcoxon rank-sum test and visualized using box plots with individual patient values. The same analysis was then extended to six melanoma anti-PD-1 cohorts from Cancer-Immu, including Auslander, Gide, Hugo, Liu, Prat, and Riaz, using only pre-treatment anti-PD-1 tumor samples. The corresponding accessions were GSE115821 (37), PRJEB23709 (38), GSE78220 (36), phs000452.v3.p1 (69), GSE93157 (70), and GSE91061 (71), respectively. TCGA-SKCM RNA-seq data were used to compare between primary melanoma and metastatic melanoma samples. Expression values were analyzed as “log2(TPM + 1)” and standardized as z-scores. The primary-vs-metastatic analysis included 103 primary tumor samples and 368 metastatic tumor samples. Group differences were tested using the Wilcoxon rank-sum test, and logistic regression was used to estimate the odds ratio for metastatic tumor status per 1-SD increase in gene or signature score. P values were adjusted using Benjamini-Hochberg FDR correction.

### Spatial processing

Processing of samples: Formalin fixed paraffin embedded Tissue Micro Array (TMA) were used. Mouse samples were profiled using in situ Xenium platform (10x Genomics). Samples were mounted onto the Xenium slides, followed by fixation of the samples according to the manufacturer’s protocol (Xenium in Situ - FFPE Tissue Preparation CG000578 | Rev F). Target transcripts were hybridized with Xenium Mouse Tissue Atlassing predesigned Gene expression probes (Xenium in Situ Gene Expression CG000749 | Rev B). After probe hybridization, ligation and amplification were performed, and the tissues were stained with cell segmentation markers overnight, followed by nuclear staining (DAPI). The slides were then loaded onto the Xenium Analyzer. Following imaging, the slides were stained for H&E (Xenium Post-Xenium Analyzer H&E Staining CG000613 | B).

TMA core extraction and cell segmentation: Whole-slide DAPI images were loaded into QuPath (v0.6.0), where rectangular regions of interest (ROIs) were manually annotated around each TMA core. Core boundary coordinates (in microns) were exported using a custom Groovy script that converts pixel coordinates to physical units using the slide’s pixel calibration. Raw Xenium transcript tables were spatially subset per core using these coordinate boundaries. Transcripts with quality value (QV) < 20 were excluded, along with BLANK, NegControl, and UnassignedCodeword probes, yielding 183.7 million quality-filtered transcripts across the 14 cores. Cell segmentation was performed using Baysor(v0.7.1) on per-core transcript CSV files with Xenium-assigned cell IDs provided as prior segmentation labels.

Quality control and processing: Baysor outputs were loaded and concatenated into a single AnnData object. Quality filtering removed cells with fewer than 5 transcripts or fewer than 3 unique genes, retaining 504,862 cells across 379 genes from an initial 506,105 Baysor-segmented cells (1,243 cells removed). Raw transcript counts were stored in a separate layer, and expression values were normalized to counts per 10,000 followed by log1p transformation.

Batch integration and dimensionality reduction with scVI: Batch effects across slides were corrected using scVI (scvi-tools v1.3.0). A variational autoencoder was trained with the following parameters: 10 latent dimensions, 128 hidden units, 1 encoder/decoder layer, zero-inflated negative binomial (ZINB) gene likelihood, maximum KL divergence weight of 1.0, and FP16 (mixed-precision) with batch size of 2048. The 10-dimensional scVI latent representation was used for all downstream analyses.

Unsupervised clustering and cell type annotation: A k-nearest neighbor graph (k = 15) was constructed on the scVI latent space using rapids_singlecell (v0.12.1). UMAP visualization, Leiden clustering, and marker gene identification were performed using scanpy (v1.10.4). UMAP embeddings were generated with min_dist = 0.1, spread = 1.0, and spectral initialization. Community detection used the Leiden algorithm at resolution 0.5. Marker genes were identified via Wilcoxon rank-sum test, retaining genes with adjusted p-value < 0.05 and >30% non-zero expression in the test group to guide cell type annotation.

Iterative subclustering and doublet removal: Each major cell type compartment was independently subclustered using the same scVI→neighbors→UMAP→Leiden pipeline (identical hyperparameters as above). Iterative rounds of sub clustering were performed per compartment with doublets, multiplets, and contaminating populations identified by aberrant co-expression of lineage-specific markers and removed at each round.

Spatial niche identification: Spatial cellular niches were identified using Cell Charter (v0.3.5). A spatial neighbor graph was constructed for each TMA core using 10 nearest neighbors based on cell centroid coordinates. Neighborhood representations were computed by aggregating scVI latent embeddings over 1-hop and 2-hop spatial neighbors. The optimal number of niches was determined using Cell Charter’s ClusterAutoK module, which evaluates cluster stability across 20 independent runs for each k in the range 5–20.

Calculation of TMA core areas: Images were imported into QuPath for analysis using standard Hematoxylin and Eosin (H&E) stain vectors. Individual TMA cores were identified and labeled using the automated TMA dearraying tool. Parameters for core detection included a core diameter of 2.0mm, a density threshold of 5.0, and a bounds scale factor of 105.0. Following automated detection, core boundaries were manually refined where necessary. To calculate the precise tissue area within each core, a machine learning-based pixel classifier was developed. Representative regions of tissue, background, and border were manually annotated across multiple cores to create a robust training set. An artificial neural network (ANN_MLP) classifier was utilized, trained, and applied at a pixel size of 4.0μm/px. The trained classifier was applied to all detected cores. The total area for each core expressed in μm^2^ was automatically calculated based on the calibrated pixel size within the image metadata. Data were exported using the “Measurement Maps” feature for downstream analysis.

Differential cell density analysis: To test for differences in cell type, subtype and spatial niche density between treatment groups, we used edgeR’s (v4.4.0) generalized linear model (GLM) quasi-likelihood (QL) framework. Cell type counts per core were organized into a count matrix (cell types x cores). Tissue area per core (mm^2^) was used as the library size (equivalent to a log-area offset), allowing direct modeling of cell densities. A design matrix encoding treatment group was specified with Ctrl as the reference level. Dispersions were estimated via glmQLFit with robust = TRUE. Differential density was assessed using the QL F-test. P-values were corrected for multiple testing using the Benjamini–Hochberg (BH) method.

Compositional analysis: Compositional differences in cell type, subtype and spatial niche proportions between treatment groups were assessed using sccomp(v2.1.17) using the sum-constrained Beta-Binomial model. The model was specified with formula ∼ 0 + group, and the contrast groupCtrl - groupIndole was tested. Outlier cells were identified and removed using sccomp_remove_outliers. Posterior probabilities of the null hypothesis (c_pH0) and BH-corrected false discovery rates were computed for each comparison.

### Statistics and Reproducibility

Statistical significance was assessed by ordinary one-way ANOVA and two-way ANOVA. GraphPad Prism 9 software (GraphPad, La Jolla, CA) performed all statistical calculations. All cell culture experiments were performed three times. Data is presented as mean ± SD (unless noted otherwise in figure legends). A *P* value <0.05 was considered statistically significant.

### Primer information

ATF4: Forward, 5’-ATGACCGAAATGAGCTTCCTG-3’; Reverse, 5’-GCTGGAGAACCCATGAGGT-3’ XBP1: Forward, 5’-AGAAGTCCGTCCACTCTCTCA-3’; Reverse, 5’-GCTCGGGATTTTACCAGTTCAT-3’ DDIT3: Forward, 5’-GGAAACAGAGTGGTCATTCCC-3’; Reverse, 5’-CTGCTTGAGCCGTTCATTCTC-3’ HSPA5: Forward: 5’-CATCACGCCGTCCTATGTCG-3’; Reverse, 5’-CGTCAAAGACCGTGTTCTCG-3’ ERN1: Forward: 5’-AGAGAAGCAGCAGACTTTGTC-3’; Reverse, 5’-GTTTTGGTGTCGTACATGGTGA-3’ PPP1R15A: Forward, 5’-ATGATGGCATGTATGGTGAGC-3’; Reverse, 5’-AACCTTGCAGTGTCCTTATCAG-3’ EIF2AK3: Forward, 5’-GGAAACGAGAGCCGGATTTATT-3’; Reverse, 5’-ACTATGTCCATTATGGCAGCTTC-3’ CYP1A1: Forward, 5’-TCGGCCACGGAGTTTCTTC-3’; Reverse: 5’-GGTCAGCATGTGCCCAATCA-3’ HERPUD1: Forward, 5’-TGCTGGTTCTAATCGGGGACA-3’; Reverse, 5’-CCAGGGGAAGAAAGGTTCCG-3’ HMOX1; Forward, 5’-AAGACTGCGTTCCTGCTCAAC-3’; Reverse, 5’-AAAGCCCTACAGCAACTGTCG-3’ JUN: Forward, 5’-TCCAAGTGCCGAAAAAGGAAG-3’; Reverse, 5’-CGAGTTCTGAGCTTTCAAGGT-3’ TGFB2: Forward, 5’-CAGCACACTCGATATGGACCA-3’; Reverse, 5’-CCTCGGGCTCAGGATAGTCT-3’ PPP2R2B: Forward, 5’-CCACACGGGAGAATTACTAGCG-3’; Reverse, 5’-TGTATTCACCCCTACGATGAACC-3’ WNT7B: Forward, 5’-GAAGCAGGGCTACTACAACCA-3’; Reverse, 5’-CGGCCTCATTGTTATGCAGGT-3’ IL1B: Forward, 5’-ATGATGGCTTATTACAGTGGCAA-3’; Reverse, 5’-GTCGGAGATTCGTAGCTGGA-3’ ATF3: Forward, 5’-CCTCTGCGCTGGAATCAGTC-3’; Reverse, 5’-TTCTTTCTCGTCGCCTCTTTTT-3’

## Data Availability

Correspondence and requests for materials should be addressed to Ze’ev A. Ronai, Jim and Eleanor Randall Departament of Surgery, the Department of Biomedical Sciences and the Translational Research Institute, Cedars Sinai Medical Center, 8700 Beverly Blvd. Los Angeles, CA, 90048, T-310-423-4248, E –

## Acknowledgments

We thank the SBP and CSMC Vivarium for helping with animal studies, the SBP Genomics core for assistance with RNA-Seq, Ramachandran Murali and Avradip Chatterjee, and Chi-Chang Yang for helping with the analysis of MA3 mutant clones. NIH support through grant R35 CA283706 and a grant from the Melanoma Research Foundation (to ZAR) are gratefully acknowledged. NCI grant P30 CA030199 enabled the contributions of shared resources to different aspects of this study. I.T. is a Senior Research Scholar of the Fonds de Recherche du Quebec-Sante (REQ-S).

## Author contributions

YF, NS, IT, HK, and ZAR designed the studies; YF, XDO, XK, MJ, and HJ performed the experiments. RM, HK, SRD, TH, and ER carried out bioinformatic analyses. MR and AC performed *in silico* screens. SO, ES, and MJ carried out small molecule screens and medicinal chemistry for select compounds. AD generated the eIF4G1 CRISPR library, and IL performed a screen for SBI-756 resistance in yeast. MA, CD, and NS carried out ribo-sequencing. IT and PJ carried out polysome profiling. JV provided MEK-resistant melanoma cells. AM performed a pathological analysis of melanoma metastasis. NS, IT, AC, CF and ZAR provided critical data assessments. ZAR, IT, MA, NS, AC, XDO, and SO wrote the manuscript.

## Abbreviations

4E-BP: Eukaryotic translation initiation factor 4E-binding protein
BCL-2 inhibitor: B-cell lymphoma 2 inhibitor
CCLE: Cancer Cell Line Encyclopedia
CDKN1A: Cyclin-dependent kinase inhibitor 1A
eIF4A: Eukaryotic initiation factor 4A
eIF4E: Eukaryotic initiation factor 4F
eIF4G1: Eukaryotic translation initiation factor 4 gamma 1
FACS: Fluorescence-activated cell sorting
FTH1: Ferritin heavy chain 1
FTL: Ferritin light chain
GO: Gene ontology
GSEA: Gene set enrichment analysis
IL24: Interleukine-24
IPA: Ingenuity pathway analysis
mTORC1: Mammalian target of rapamycin complex 1
NRF2: Nuclear factor erythroid 2-related factor 2
PCA: Principal component analysis
p53: Encoded by the TP53 gene
PD-1: Programmed cell death protein 1
ROS: Reactive oxygen species
TGFβI: Transforming growth factor beta-induced protein
THBS1: Thrombospondin-1
TIF1: transcriptional intermediary factor 1
UPR: Unfolded protein response

## Supplementary Figure Legends

**Supplementary Figure 1.**
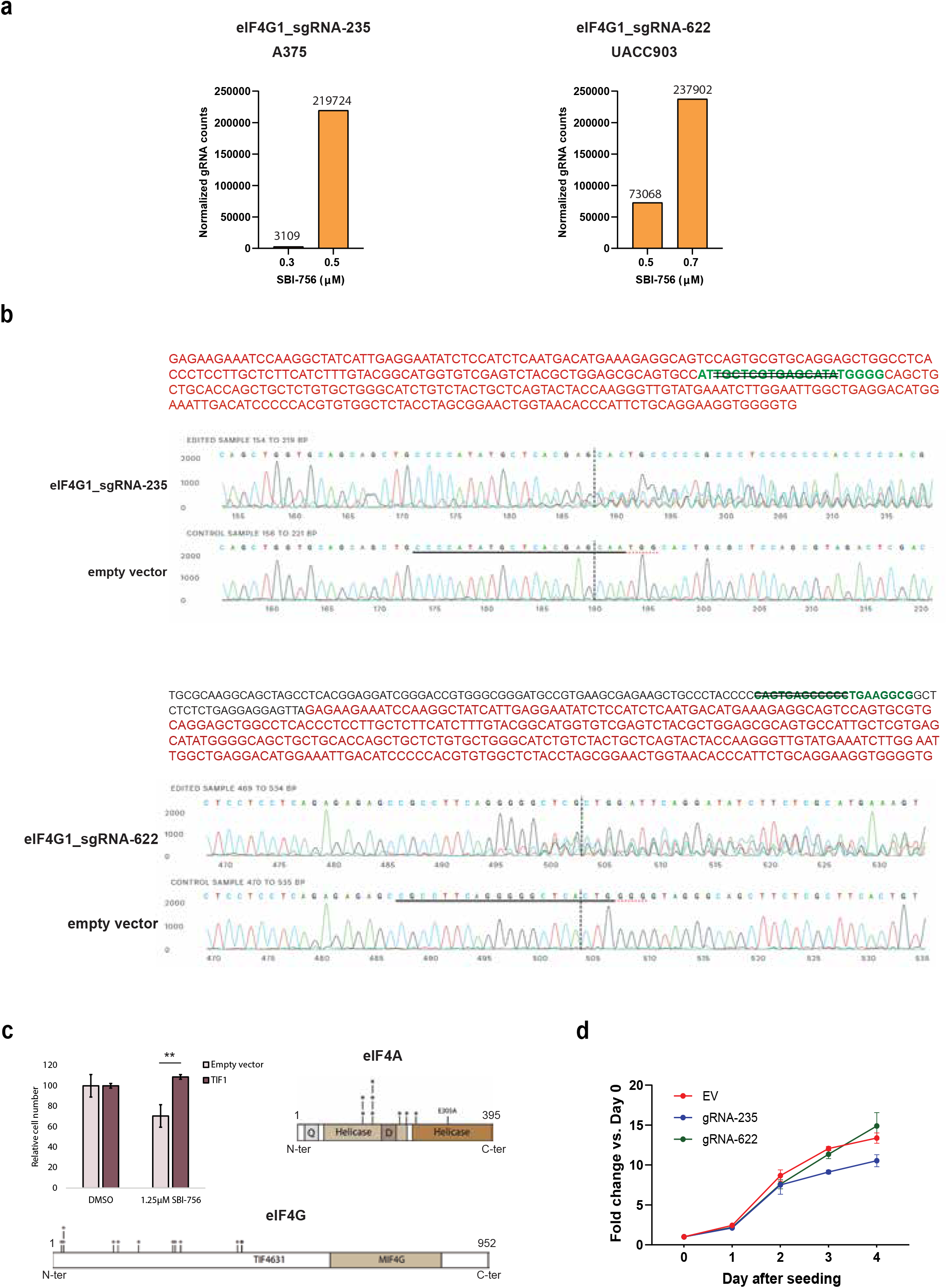
Identifying MA3 domain as SBI-756 target on eIF4G1. **(a)** A375 and UACC903 melanoma cells were subjected to indicated sgRNA infection followed by treatment with SBI-756. Genomic DNA was extracted from the cells followed by sequencing. eIF4G1_sgRNA-235 and sgRNA-622 counts were shown. **(b)** MA3 domain sequences of gRNA-235 and gRNA-622 A375 cells. sgRNA sequence shown in green and MA3 sequence in red. **(c)** Left, KY775 strain was transformed with an empty vector or a vector coding for TIF1. Transformed cells were grown with DMSO or 1.25μM SBI-756 for 48h before assessing the cell number by optical density. Data were normalized to DMSO treated cells. Error bars represent SEM, n=3. Significance was assessed by Student’s T test: **: p-value < 0. Right, KY775 strain was transformed with a library of randomly mutated TIF1 expressing vectors. 9 clones resistant to SBI-756 were retrieved and DNA was extracted, transformed into TOP10 competent bacteria. Plasmids present in each clone were sequenced. All clones presented deletion or mutation of the helicase C-terminal domain. Stars indicate the presence of a stop codon. Q: Q motif, D: DEAD box. Bottom, TIF4631 deletions promote KY775 *S. cerevisiae* (pdr5D::hisG erg6D::hisG) strain resistance to SBI-756. KY775 cells were transformed with a library of randomly mutated TIF4631 expressing vectors. 12 clones resistant to SBI-756 were retrieved and DNA was extracted, transformed into TOP10 competent bacteria. Plasmids present in each clone were sequenced. All clones presented deletion of the C-terminal domains of TIF4631. Stars indicate the presence of a stop codon. MIF4G: middle domain of eukaryotic initiation factor 4G. **(d)** gRNA-235, gRNA-622 and EV A375 cells were seeded in 96-well plates. Cell proliferation was measured daily for 4 days using CellTiter-Glo.

**Supplementary Figure 2.**
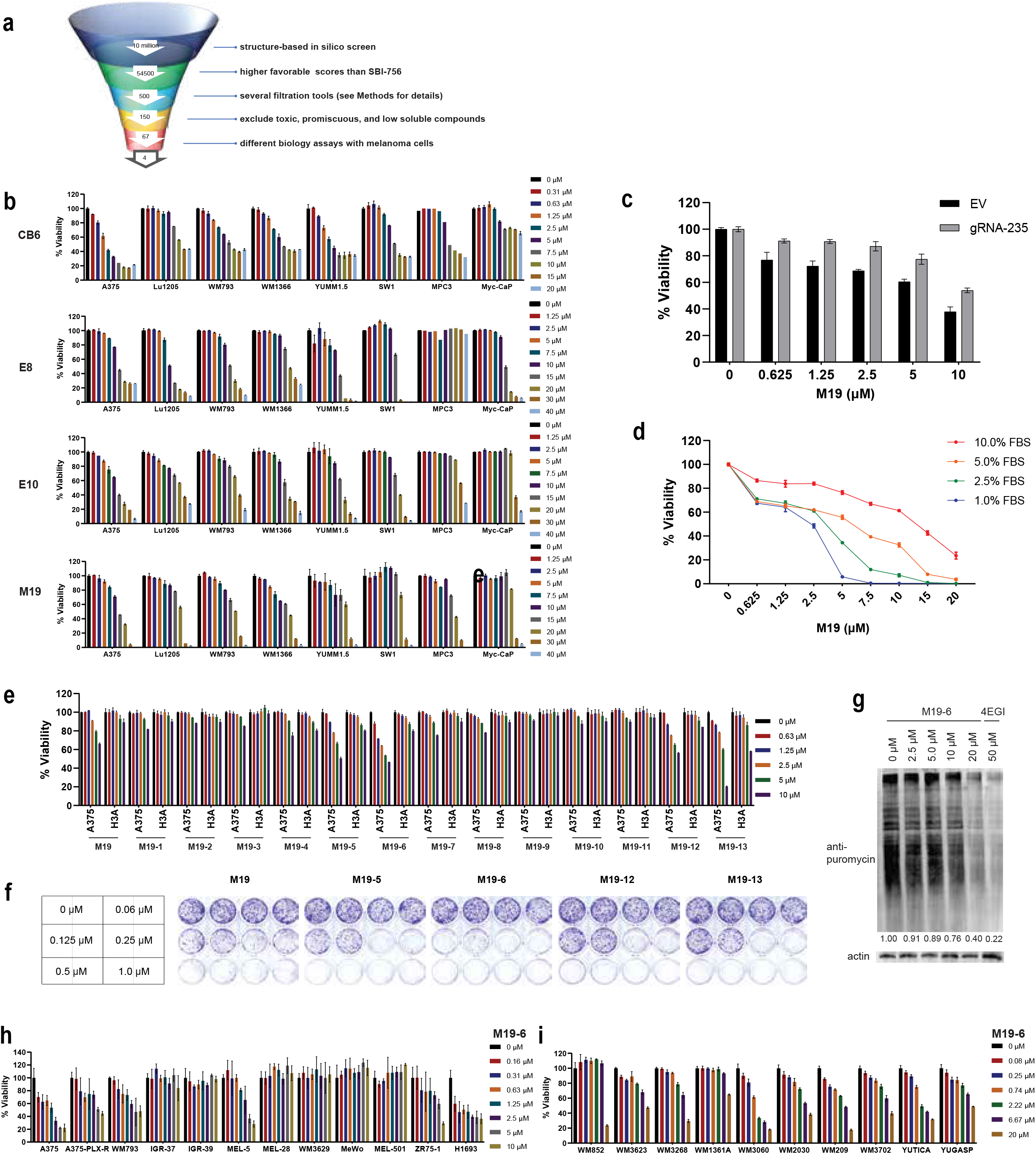
Characterizing MA3 domain bound small molecules selected from in Silico screen. **(a)** Testing funnel of high-throughput screening campaign (details in Methods). **(b)** Indicated melanoma and prostate cancer cell lines were treated with the four selected compounds, followed by viability assay after 72h. **(c)** EV and gRNA-235 A375 cells were treated with M19, followed by viability assay. **(d)** A375 cells were cultured in a medium containing indicated concentrations of FBS, followed by treatment with M19. Cell viability was measured 72 h later. **(e)** A375 melanoma and H3A melanocyte cells were treated with indicated M19 analogs. Cell viability was measured 72 h later. **(f)** A375 cells (300 cells/well) were seeded in 12-well plates and treated with indicated compounds. After one week, cells were stained with crystal violet. **(g)** A375 cells were pre-incubated with indicated concentrations of M19-6 for 1 h, followed by incubation with 3 μg/mL puromycin for 0.5 h. Cells were collected, lysed, and subjected to Western blot analysis using anti-puromycin antibody. The density of the whole lane was quantified by ImageJ and normalized to actin levels as indicated. **(h)** Indicated BRAF mutant melanoma and breast cancer cells grown in 2D cultures or as spheroids were treated with M19-6, and cell viability was measured (after 72 h for cell cultures and 96 h for spheroids). **(i)** Ten different NRAS-mutant melanoma cell lines (seeded in 384-well plates) were treated with the indicated concentrations of M19-6. Cell viability was measured after 72 h.

**Supplementary Figure 3.**
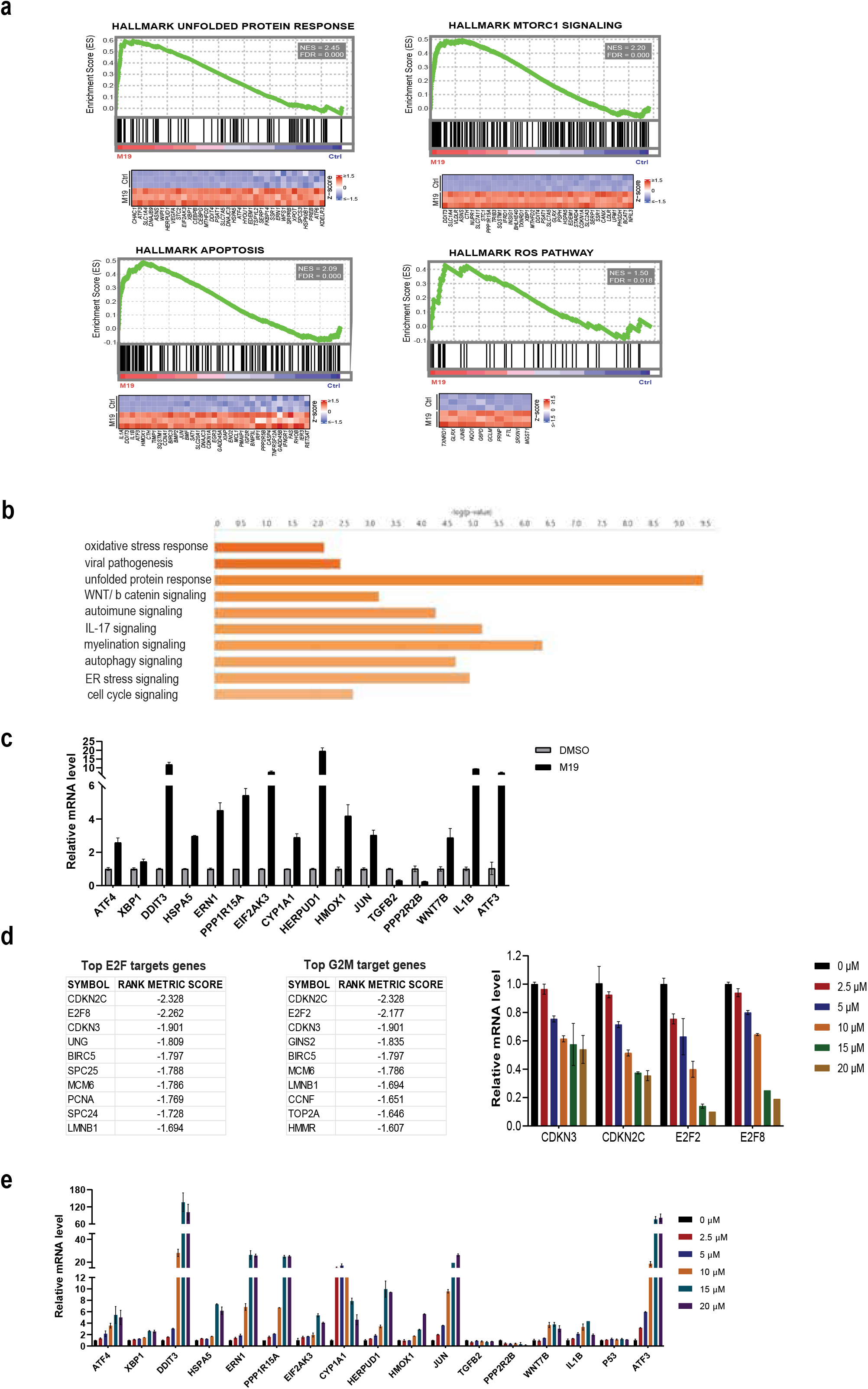
RNA-Seq analysis of A375 cells treated with M19. **(a)** Gene set enrichment analysis of RNA-Seq data of A375 cells treated with M19 (20 μM). The top enrichment plots of indicated pathways and corresponding heatmaps of up-regulated genes are shown. **(b)** IPA analysis of RNA-Seq data of A375 cells treated with M19 (20 μM); top activated pathways are shown. **(c)** qPCR validation of selected differentially expressed genes. **(d)** Representative genes and related signaling that were downregulated upon M19 treatment (left panel) and their validation by qPCR (right panel). **(e)** A375 cells were treated with the indicated concentrations of M19-6. qPCR analysis was performed on the indicated, differentially expressed, genes.

**Supplementary Figure 4.**
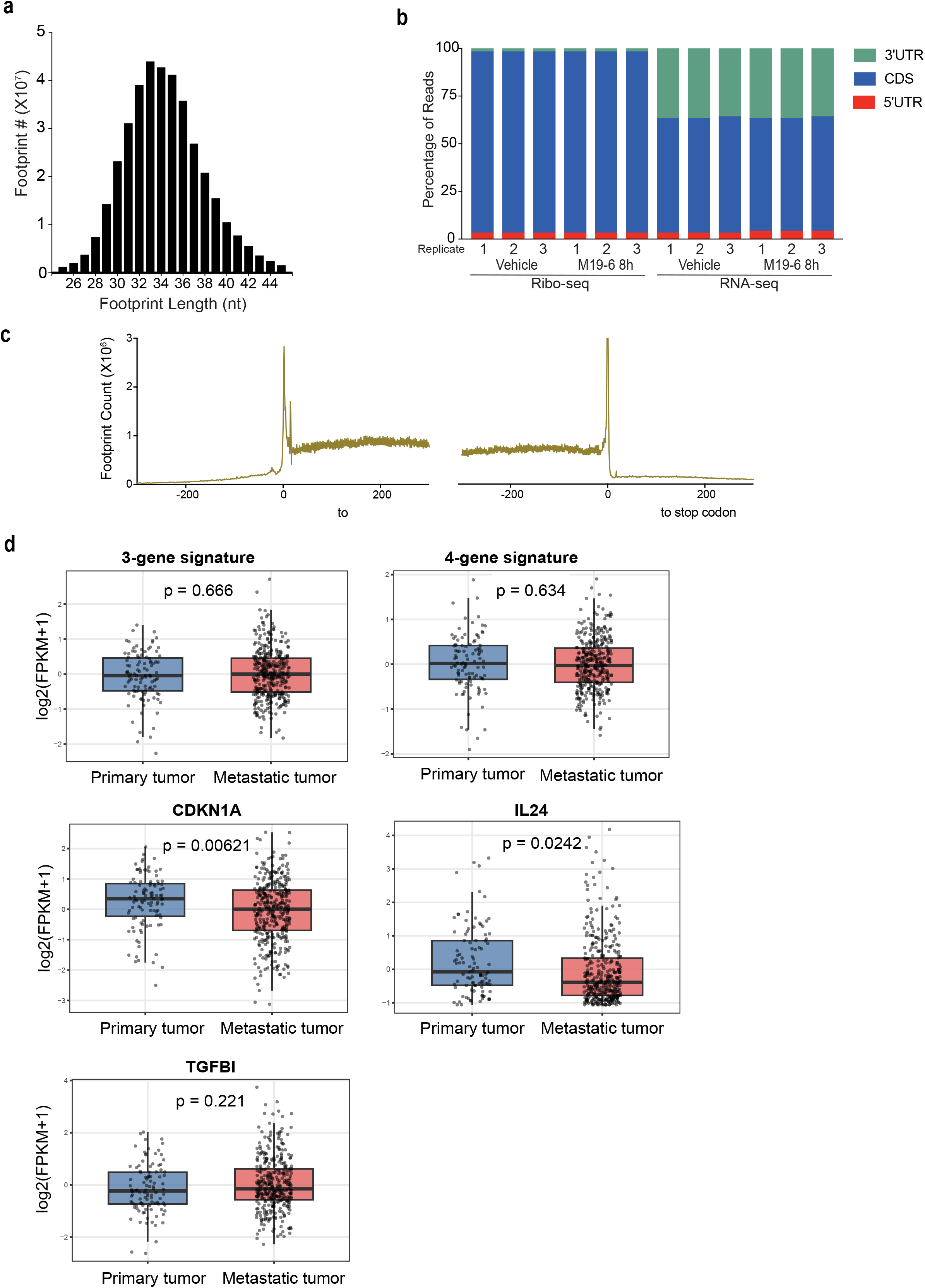
Characterization of ribosome footprints. **(a)** Distribution of read lengths for ribosome footprints generated using MNase digestion. **(b)** Stacked bar charts showing the proportion of reads from RNA-Seq and Ribo-Seq libraries mapped to different mRNA regions: coding sequence (CDS), 5’ untranslated region (5’ UTR), and 3’ untranslated region (3’ UTR). **(c)** Metagene analysis of ribosome profiling libraries generated by MNase, with reads from all mRNAs aligned within ±300 nucleotides of the start and stop codons. **(d)** TCGA-SKCM RNA-Seq data were used to compare expression of TGFβI, IL24, CDKN1A, and the indicated combined signatures between primary and metastatic melanoma samples.

**Supplementary Figure 5.**
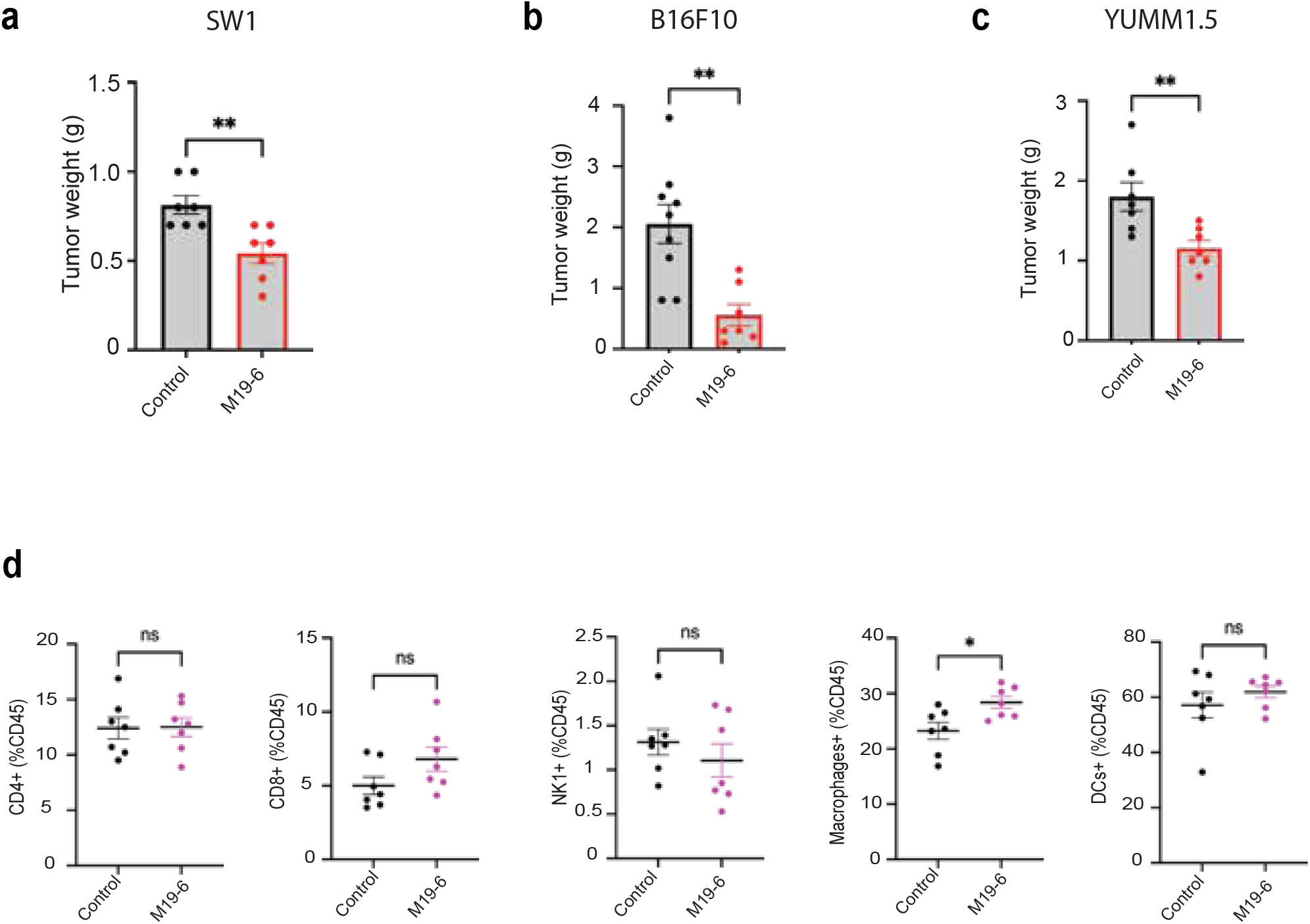
M19-6 inhibits melanoma growth by anti-tumor immunity. **(a)** SW1 cells grafted into 8-week-old C3H/HeNHsd mice (n=10 per experimental group) subjected to treatment with M19-6. Shown is the weight of SW1 tumors. **(b)** B16F10 cells were inoculated in C57BL/6 mice (n=10 per experimental group) that were subjected to treatment with M19-6. Shown is the weight of B16F10 tumors. **(c)** YUMM1.5 cells were inoculated in C57BL/6 mice (n=10 per experimental group) for 7-10 days before emerging tumors were subjected to treatment with M19-6. Weight of the tumors at indicated time points is shown. **(d)** Quantification of tumor infiltration of immune cells (CD8^+^ T cells, CD4^+^ T cells, NK1, DCs and macrophages) 12 days after tumor inoculation of YUMM1.5 cells into C57BL/6J mice that were treated with M19-6 or vehicle (n=7 mice per treatment). Data shown represent two experiments. Data were analyzed by unpaired t-test. *P < 0.05, **P < 0.005, ***P < 0.001, ****P < 0.0001 by two-tailed t-test or two-way ANOVA.

